# Cellular nucleic acid-binding protein (CNBP) dependent cytokine programming shapes host defense against *Plasmodium* infection

**DOI:** 10.64898/2026.06.01.729202

**Authors:** Romana Rashid, Leandro de Souza Silva, Shahid Banday, Daniel R Caffrey, Evelyn A Kurt-Jones, Katherine A Fitzgerald, Douglas T Golenbock

## Abstract

During blood-stage *Plasmodium* infection, effective immune control hinges on the IL12β-IFN-γ axis, yet how this pathway is transcriptionally tuned *in vivo* remains incompletely defined. Innate sensing of parasite-derived ligands by pattern-recognition receptors, including Toll like receptors, in dendritic cells and macrophages induces IL-12β production that drives IFN-γ mediated control of infection. Emerging evidence implicates cellular nucleic acid-binding protein (CNBP), a zinc-finger transcriptional regulator, in control of IL12β gene expression in myeloid cells exposed to bacterial and viral infections.

Here, we defined the contribution of CNBP in cytokine-driven immunity to *Plasmodium* infection including both *P. falciparum* (the major cause of malaria), as well as *P. chabaudi* and *P. berghei* ANKA, two rodent species that model human disease. Upon exposure to *Plasmodium*-infected erythrocytes, CNBP rapidly translocated to the nucleus in mouse and human dendritic cells, bound IL12β promoter, and was required for optimal IL12β induction. Genetic ablation of CNBP in mice and siRNA knockdown of CNBP in human monocyte-derived dendritic cells markedly reduced IL12β production and downstream IFN-γ responses, while TNF-α and several other innate cytokines were largely unaffected. *In vivo*, hematopoietic-specific deletion of CNBP (using *vav-iCre; Cnbp^fl^*^/fl^) resulted in elevated peak parasitemia, impaired parasite clearance, and relapse after initial resolution. Consistent with these outcomes, spleens from mice lacking CNBP in hematopoietic cells exhibited reduced inflammatory remodeling, altered T-cell composition, and transcriptional reprogramming characterized by selective regulation of IL12β-IFN-γ transcripts alongside upregulation of distinct cytotoxic genes. Paradoxically, mice lacking CNBP in hematopoietic cells showed delayed mortality in the lethal infection model, underscoring its context-dependent contributions to host protection and inflammatory pathology.

Collectively, these findings position CNBP as a pivotal modulator of the IL12β-IFN-γ axis during malaria, extending its functional repertoire beyond microbial contexts, with potential as a therapeutic target to fine-tune immune responses for enhanced protection with limited immunopathology.

**Author summary:** Malaria remains one of the world’s deadliest infectious diseases, caused by the protozoan *Plasmodium* that trigger complex immune responses in infected hosts. Effective host defense requires a tightly regulated inflammatory response: too weak and the parasite proliferates unchecked; too strong and the host suffers harmful immunopathology. Central to this balance is the IL12β-IFN-γ signaling axis. In this study, we investigated the role of a transcriptional regulator, known as Cellular Nucleic acid-Binding Protein (CNBP), in shaping the immune response against the *Plasmodium* parasite. We found that CNBP rapidly responds to parasite exposure, translocates to the nucleus and drives IL12β production in innate immune cells, and promotes downstream IFN-γ driven adaptive immune responses. Mice lacking CNBP in hematopoietic stem cells exhibited changes in transcription of key inflammatory genes; markedly reduced systemic IL12β and IFN-γ, leading to significantly elevated peripheral blood parasitemia and altered immune cell composition. Paradoxically, the absence of CNBP delayed mortality in a lethal infection model and reduced inflammatory responses that are associated with cytokine storm-mediated immunopathology. These findings identify CNBP as a key regulator that fine-tunes protective immunity and inflammatory pathology during malaria, highlighting its potential as a therapeutic target to optimize host defense while limiting harmful inflammation.

## Introduction

*Plasmodium spp*., the causative agents of malaria, remain a major global health threat, responsible for over 610, 000 deaths in 2024 alone. Young children in sub-Saharan Africa carry the heaviest burden **[1]**. Effective host resistance to *Plasmodium* critically depends on the early activation of innate immune responses and the induction of pro-inflammatory cytokines **[2]**. A central component of this defense is interleukin-12 (IL-12), produced by dendritic cells and macrophages, which drives Th1 differentiation and stimulates CD4⁺ and CD8⁺ T cells as well as NK cells to produce interferon-γ (IFN-γ). IFN-γ in turn activates macrophages, enhancing parasite killing **[3,4,5]**. However, excessive cytokine production can result in systemic inflammation and contribute to severe malaria, in particular cerebral malaria, life-threatening anemia or metabolic acidosis **[6,7,8]**. Thus, a fine balance between pro-inflammatory mediators, IL-12 and IFN-γ, and regulatory cytokines like IL-10, is essential to control parasites while limiting host damage **[9]**.

Innate immunity is the first line of defense against Plasmodium. During blood-stage infection, parasite components from infected erythrocytes are sensed by pattern-recognition receptors on macrophages, dendritic cells, and other innate cells **[10]**. Parasite glycosylphosphatidylinositol (GPI) anchors activate macrophages to produce TNF-α and IL-12 via Toll-like receptor pathways **[11]**. Parasitic DNA associated with hemozoin stimulates nucleic-acid sensing pathways in dendritic cells, promoting IL-12 production and Th1 responses **[12,13,14]**. Subsequent work demonstrated that AT-rich DNA motifs from the Plasmodium genome activate innate immune pathways and type I interferon responses, highlighting the importance of nucleic-acid sensing during malaria infection **[15]**. Collectively, innate immune recognition of Plasmodium-derived ligands by macrophages and dendritic cells initiates inflammatory cytokine programs that shape the development of protective Th1 immunity during malaria [**4,16].**

Recent work has underscored the importance of transcriptional and post-transcriptional mechanisms in tuning innate immune responses. RNA-binding proteins and nucleic acid-interacting factors have emerged as key regulators of cytokine gene expression by influencing mRNA stability, translation, and transcription factor activity **[17,18,19,20]**. These discoveries have opened new avenues for exploring how cytokine responses are fine-tuned during infections like malaria. One such regulator is the Cellular Nucleic Acid Binding Protein (CNBP), a conserved zinc finger protein, initially characterized for nucleic acid binding and RNA metabolism [**21]**. CNBP has more recently been implicated as a transcriptional regulator in innate immune cells. CNBP has been shown to regulate the NF-κB family member c-Rel, which is uniquely required for IL-12β gene transcription, and CNBP-deficient mice exhibit defective IL-12β induction and impaired Th1 responses, leading to increased susceptibility to intracellular pathogens such as *Toxoplasma gondii* **[22,23]**. In addition, CNBP contributes to sustained IL-6 production **[24]** and has been reported to regulate type I interferon responses via interactions with the interferon regulatory factors (IRFs), IRF3 and IRF7 **[25]**. These studies have collectively positioned CNBP as a gatekeeper of cytokine-driven immunity and raise the possibility that CNBP may regulate the immune response during *Plasmodium* infection.

Despite these insights, the role of CNBP in many protozoan infections, including malaria, remains largely unexplored. This gap is particularly significant, given the central role of the IL12-IFN-γ signaling axis in host resistance to *Plasmodium* **[26,27,28]**. Whether CNBP regulates this axis during malaria infection remains an unexplored area. Here, we investigated the activation and contribution of CNBP to cytokine programming and host defense in murine models of *Plasmodium* infection, complemented by studies in human monocyte-derived dendritic cells.

We employed a hematopoietic cell - specific CNBP knockout mouse model developed by crossing CNBP floxed alleles to Vavi-Cre delete strains and examined host responses in both the self-resolving *Plasmodium chabaudi* AS model and the lethal *Plasmodium berghei* ANKA model. We assessed parasitemia, survival, cytokine production, and splenic immune architecture, and complemented these studies with analyses of CNBP localization in human monocyte-derived dendritic cells stimulated with *Plasmodium falciparum*-infected red blood cells.

Our findings demonstrate that CNBP is required for optimal induction of the IL12-IFN-γ axis and for effective parasite control, while also influencing the magnitude of inflammatory responses that contribute to disease outcome. These results identify CNBP as a key regulator of cytokine-driven immunity during malaria infection and provide new insight into the transcriptional networks that balance protective host defense with immunopathology in protozoan infection.

## Results

### CNBP translocates to the nucleus of dendritic cells exposed to *Plasmodium* infected RBCs

Despite extensive study of pattern recognition pathways in malaria, how nuclear transcriptional co-regulators respond to parasitic ligands is not well-defined. CNBP (Cellular nucleic acid binding protein), previously implicated in bacterial and viral immunity via *Il12b* regulation [**22,24,25,29]**, has not been investigated in the context of *Plasmodium* parasites.

To examine CNBP localization in response to parasite-derived stimuli, bone marrow-derived dendritic cells (BMDCs) from wild-type C57BL/6 mice were exposed to *P. falciparum*-infected erythrocytes (iRBCs) *in vitro* for up to 24 h, with the 0 h time point representing mock-treated cells. Subcellular fractionation followed by immunoblotting revealed marked CNBP enrichment in the nuclear fraction of iRBC-stimulated BMDCs, while mock treated cells did not exhibit this translocation. The purity of nuclear and cytosolic fractions was confirmed using USF2 and GAPDH antibody, respectively (**Fig 1A**). We next sought to determine whether the CNBP nuclear translocation response was conserved in human dendritic cells. CD14⁺ monocytes from healthy donors were differentiated into monocyte-derived dendritic cells (MoDCs) and stimulated with *P. falciparum iRBCs* (1:10 ratio) for the indicated times (0, 2, 4 and 6 h). Immunoblotting of nuclear and cytosolic fractions revealed time-dependent nuclear accumulation of CNBP at 4 and 6 h post stimulation. In contrast, stimulation with uninfected erythrocytes (uRBCs) failed to induce this translocation, closely mirroring the observation of murine BMDCs (**Fig 1B**). Lipopolysaccharide (LPS, 100 ng/ml x 30 min), a known activator of CNBP via NF-κB pathways, was included as a positive control for nuclear translocation. These findings were further corroborated by immunofluorescence microscopy in BMDCs, which revealed robust nuclear localization of CNBP after 6 h of iRBC stimulation (**Fig 1C**). Consistent with these observations, stimulation of MoDCs with natural hemozoin (100 µM) isolated from *P. falciparum* cultures also induced progressive CNBP nuclear accumulation. Subcellular fractionation and immunoblot analysis showed predominantly cytosolic CNBP at baseline, with faint nuclear localization at 2 h that increased at 4 h and became prominent by 6 h (**Fig S1A**), indicating that exposure to hemozoin alone is sufficient to drive time-dependent CNBP nuclear translocation.

**Figure 1.**
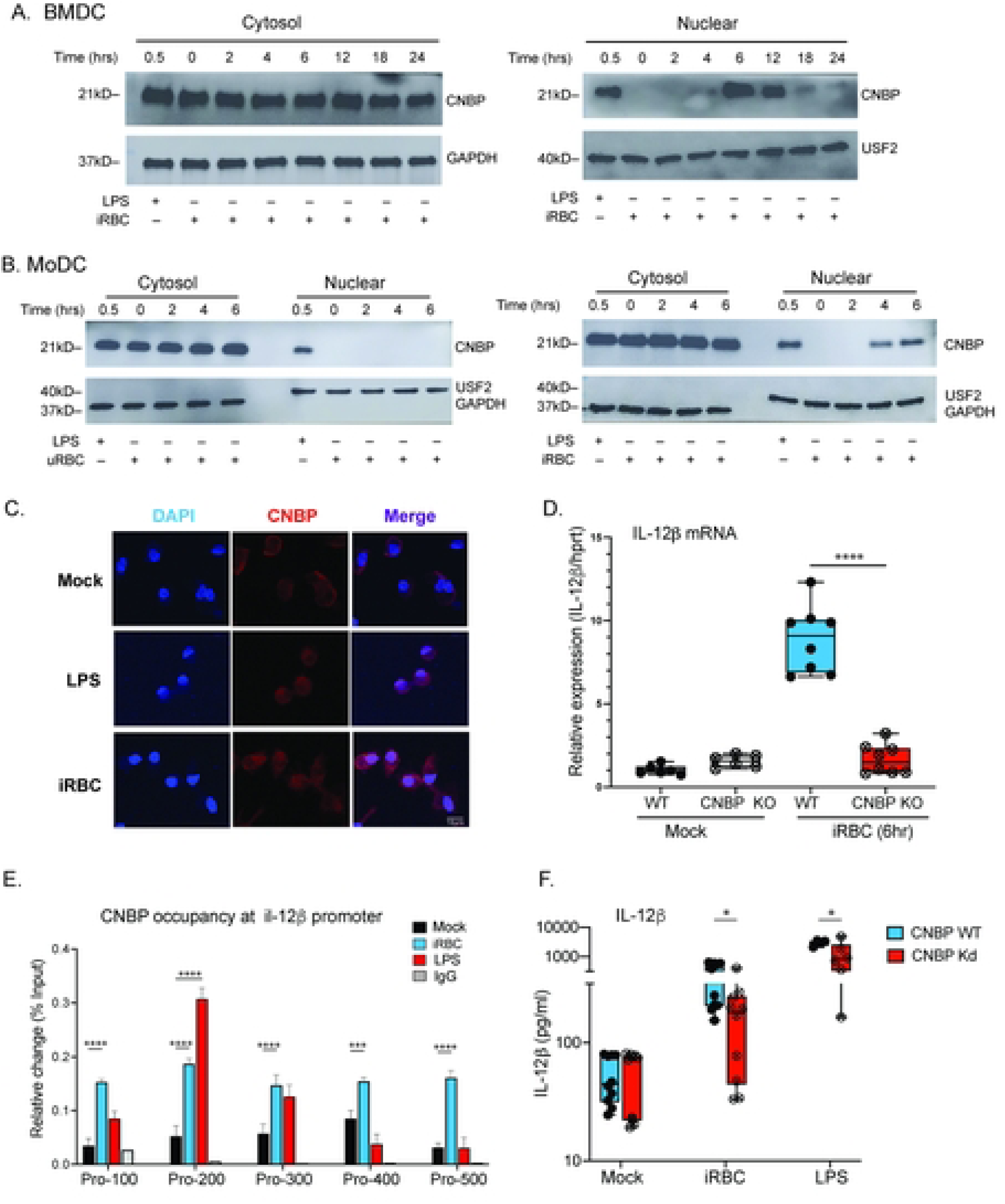
CNBP translocates to the nucleus in murine and human dendritic cells in response to *Plasmodium falciparum*-infected erythrocytes. (A) Immunoblot of nuclear and cytosolic fractions from murine bone marrow-derived dendritic cells (BMDCs) stimulated *in vitro* with synchronized trophozoite-stage *P. falciparum*-infected red blood cells (BMDC:iRBC ratio of 1:10) at the indicated times. LPS (100 ng/mL, 30 min) served as a positive control. USF2 and GAPDH were used as nuclear and cytoplasmic markers, respectively. Each lane was loaded with 15μg of protein. Representative blots from three independent experiments are shown. (B) Immunoblot of cytoplasmic and nuclear fractions from human monocyte-derived dendritic cells (MoDCs) stimulated with *P. falciparum*-iRBCs or matched uninfected RBCs (derived from the same donor used for parasite culture) for 0-6 h. LPS (100 ng/mL, 30 min) served as a positive control. Representative blots from three independent experiments are shown. (C) Immunofluorescence images of MoDCs stimulated with iRBCs (6 h) or LPS (100 ng/mL, 30 min), stained for CNBP (Alexa Fluor 647, red) and nuclei (DAPI, blue). Images were acquired using a Leica SP8 confocal microscope (63x oil objective). Representative images from one donor are shown; the experiment was performed twice. (D) *Il12b* mRNA expression in WT and CNBP-deficient BMDCs stimulated with iRBCs or mock treated for 6 h, measured by qPCR and normalized to *Hprt*. Data represent mean ± SEM from 6-8 biological replicates. Statistical significance was determined by two-way ANOVA; ****, P<0.0001. (E) Chromatin immunoprecipitation (ChIP) using anti-CNBP antibody in BMDCs stimulated with trophozoite-stage iRBCs. Enrichment at the Il12b promoter (-100 to -500 bp; primer sequences in Sup. Table S1) is shown as relative fold enrichment over input; IgG served as a control antibody. LPS (100 ng/mL, 30 min) was used as a positive control. Data are representative of three independent experiments. Statistical significance was determined by two-way ANOVA; ***, P<0.001; ****P<0.0001. (F) IL-12β secretion by MoDCs in culture supernatant following CNBP siRNA knockdown (Kd) or NTC siRNA (non-targeting control siRNA) and stimulation with iRBCs (24 h). Each point represents an individual donor. Data are pooled from two independent experiments, each including five donors (total n = 10 donors). Statistical comparisons were performed using paired two-tailed Student’s t-test with false discovery rate correction (two-stage Benjamini-Krieger-Yekutieli method); adjusted P < 0.05 was considered significant.

To determine the transcriptional consequences of this nuclear translocation, we quantified *Il12*β expression in wild-type and CNBP deficient BMDCs following iRBC stimulation. Whereas BMDCs from wildtype mice exhibited robust induction of *Il12*β mRNA, CNBP deficient cells displayed markedly reduced transcript levels, indicating that CNBP is required for optimal Il12β gene expression downstream of *Plasmodium* sensing (**Fig 1D**). Consistent with a direct transcriptional role for CNBP, we performed chromatin immunoprecipitation (ChIP) in BMDCs exposed to *Pf* iRBCs to evaluate CNBP engagement at the *Il12*β locus. CNBP occupancy increased markedly after stimulation, with prominent enrichment across multiple regions of the promoter (-100 to -500 bp upstream of the transcription start site). In contrast, unstimulated (mock) BMDCs showed negligible CNBP binding, indicating minimal basal promoter occupancy. As a positive control, LPS (30 min) induced strong CNBP enrichment at approximately -200 bp, confirming the responsiveness of this promoter region to TLR4 mediated signaling (**Fig 1E**). Together, the data reveals that CNBP is recruited to the Il12β promoter upon pathogen challenge with stimulus dependent promoter engagement in dendritic cells.

The conservation of CNBP nuclear translocation across murine and human systems, together with direct Il12β promoter engagement in murine BMDCs, prompted us to examine whether this regulatory axis translates into functional cytokine output in human innate immune cells. Therefore, we assessed IL12β secretion by utilizing MoDCs following CNBP knockdown. In human MoDCs, CNBP knockdown (CNBP *Kd*) was achieved using siRNA transfection (150 nM), with significant knockdown efficiency of protein levels (**Fig S1B**). CNBP *kd* MoDCs secreted substantially less IL12β after 24 h of *Pf*. iRBC stimulation compared with control treated cells (**Fig 1F**). Together, these results demonstrate that CNBP supports stimulus-dependent IL12β transcription and cytokine output in response to *Plasmodium* iRBCs in murine and human dendritic cells.

### CNBP deficiency is associated with higher parasitemia and altered survival kinetics

We next assessed whether this conserved regulatory activity is associated with parasite control and disease progression. To investigate the role of CNBP in *Plasmodium* infection *in vivo*, we generated C57BL/6 mice with hematopoietic cell-specific CNBP deletion (CNBP^fl/fl^ × Vav-iCre^+/+^) by crossing CNBP^fl/fl^ mice with *Vav-iCre* transgenic mice (**Fig S2A**). Littermate CNBP^fl/fl^ mice served as wild-type controls. Hereafter, CNBP^fl/fl^ × Vav-iCre^+/+^ mice are referred to as CNBP-deficient (CNBP KO), and CNBP^fl/fl^ mice as wild-type (WT).

Eight- to ten-week-old mice were infected intraperitoneally with 1×10^6^ GFP-expressing *Plasmodium chabaudi* AS-GFP or *Plasmodium berghei* ANKA-GFP infected RBCs (**Fig 2A**). Parasitemia was determined from tail vein blood using Giemsa-stained thin smears (**Fig 2B**) as well as by flow cytometric quantification of Ter119^+^ GFP^+^ erythrocytes, enabling enhanced sensitivity and dynamic range. During *P. chabaudi* infection, monitored over four weeks, CNBP KO mice exhibited significantly elevated parasitemia from day 5 post-infection, reaching higher peak levels than WT controls between days 7-9 (mean peak parasitemia ∼43% vs. ∼32%). Following initial parasite clearance, CNBP KO mice displayed a marked recrudescence between days 15-18 (**Fig 2C**). Infection associated weight loss was even more pronounced in CNBP KO mice (**Fig 2D**). As *P. chabaudi* infection is a non-lethal model of malaria, survival analysis was not applicable.

**Figure 2:**
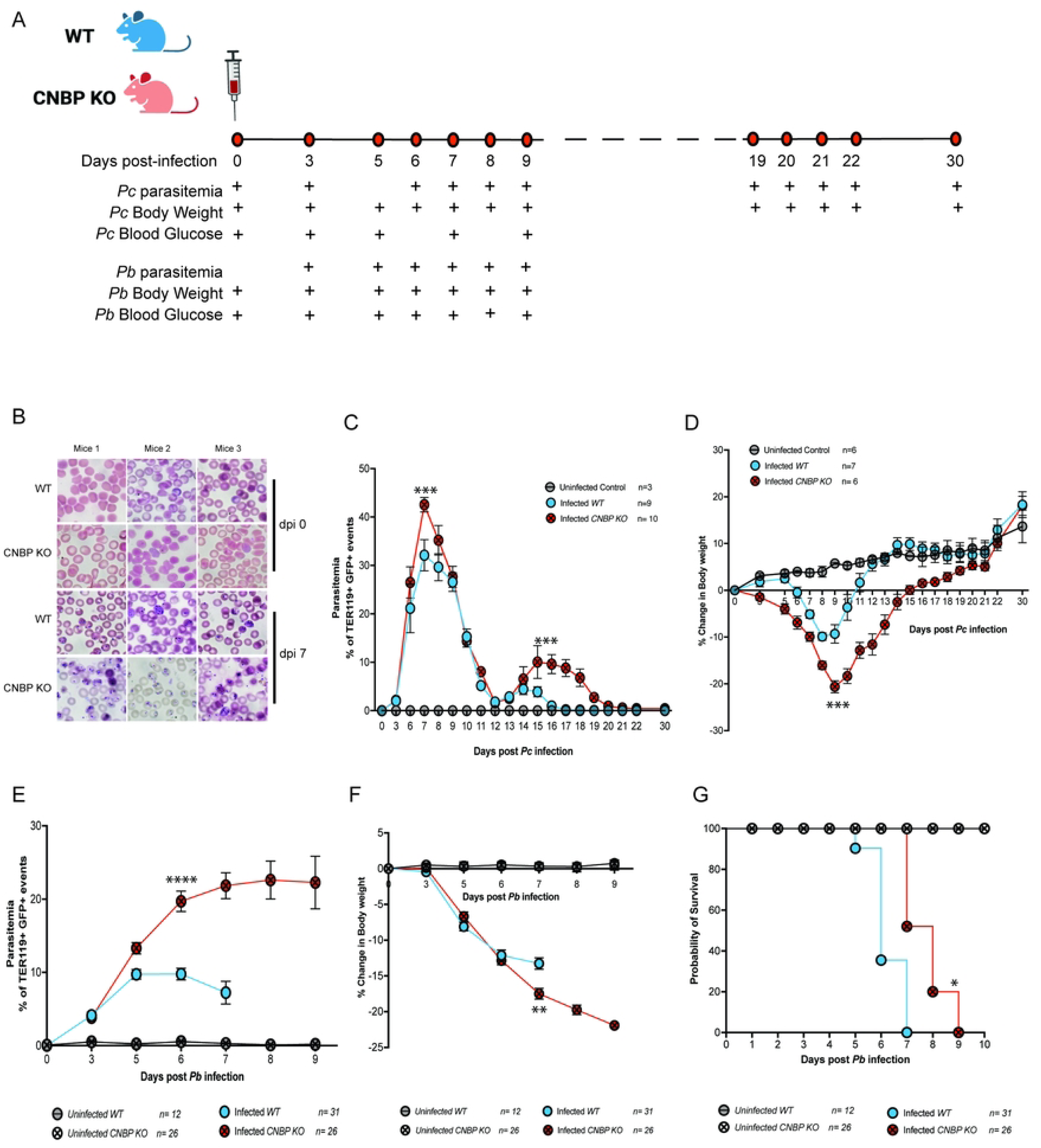
Loss of CNBP exacerbates blood-stage parasitemia and affects mortality dynamics in experimental malaria. (A) Experimental schematic illustrating intraperitoneal infection of mice with *Plasmodium* spp., followed by longitudinal monitoring of parasitemia, body weight, blood glucose, and survival at the indicated times. Peripheral blood was collected for Giemsa staining and flow cytometric analysis. (B) Mice were infected intraperitoneally with 1×10⁶ infected red blood cells (iRBCs). Representative Giemsa-stained peripheral blood smears from WT and KO mice at day 0 and day 7 PI are shown. Images are from one mouse per experimental group and are representative of three independent experiments. (C) Parasitemia kinetics in *P. chabaudi*-infected wild-type (WT; n = 11) and CNBP knockout (KO; n = 10) mice; uninfected controls (n = 3). Flow cytometric quantification of Ter119⁺GFP⁺ erythrocytes in *P. chabaudi*-infected mice demonstrating significantly higher peak parasitemia in KO mice at days 6-8 PI (day 7: p = 0.0001; day 8: *P* = 0.02) and relapse between days 15-18 PI (\*\*\**P* = 0.0099-0.0001; two-way ANOVA with multiple-comparison testing). Data are representative of one experiment from 4 independent repeats. (D) Body weight change in *P. chabaudi*-infected WT (n = 7) and KO (n = 6) mice, normalized to baseline (day 0). KO mice exhibited greater weight loss during peak parasitemia (days 8-13 PI). Statistical analysis was performed using two-way ANOVA with multiple comparisons; ***, *P* = 0.001. Data are presented as mean ± S.E.M. (E) Parasitemia in *P. berghei* ANKA-infected WT (n = 9) and KO (n = 10) mice measured by flow cytometry. KO mice reached ∼20% parasitemia at day 6 PI and maintained elevated levels (25-28%) through days 7-9 PI, whereas WT mice peaked at ∼10-12% at day 6 PI. Statistical analysis was performed using two-way ANOVA with multiple comparisons; ****, *P* = 0.0001. (F) Body weight change in *P. berghei* ANKA-infected WT (n = 31) and KO (n = 26) mice normalized to day 0. KO mice exhibited greater weight loss at peak parasitemia (day 7 PI), and weight decline continued in the surviving animals. Statistical analysis was performed using two-way ANOVA with multiple comparisons; \*\**P* = 0.003. Data are presented as mean ± SEM. (G) Kaplan-Meier survival analysis of *P. berghei* ANKA-infected WT (n = 31) and KO (n = 26) mice. WT mice succumbed between days 5-7 PI, whereas KO mice showed delayed mortality (days 7-9 PI). Survival differences were analyzed using the log-rank (Mantel-Cox) test; *P* < 0.01.

To assess the impact of CNBP deficiency under a lethal infection model, we employed *P. berghei* ANKA (*Pb* ANKA), which induces experimental cerebral malaria (ECM). Consistent with findings from *P. chabaudi* infection, CNBP KO mice developed higher parasitemia, reaching mean levels of 20-28% compared with 10-12% in WT controls during the peak acute phase of infection (**Fig 2E**). Similarly, progressive weight loss was observed in CNBP KO mice over the course of infection in the ECM model (**Fig 2F**). Notably, CNBP KO mice exhibited a modest but significant extension in median survival by approximately 1.5 days, with 50% surviving to day 7 and 20% beyond day 8, whereas all WT counterparts succumbed between days 6 and 7 (**Fig 2G**). Body weight and blood glucose levels were monitored during the acute phase as indicators of disease progression (**Fig. S2B, S2C**).

Collectively, these data demonstrate that CNBP deficiency is associated with impaired control of blood-stage parasitemia across distinct malaria models and with altered survival dynamics during experimental cerebral malaria.

### CNBP deficiency limits splenic enlargement despite enhancing hemozoin accumulation

The spleen is the primary site for clearance of infected RBCs and hemozoin during blood-stage malaria [**30]**. To examine the role of CNBP in splenic remodeling during *Plasmodium* infection *in vivo*, spleens harvested from 8-week-old CNBP KO and WT mice were analyzed at defined time points following *P. chabaudi* or *P. berghei* infection. Histological analysis of uninfected spleens demonstrated well defined red and white pulp organization in WT mice, whereas CNBP-deficient spleens exhibited subtle alterations in splenic architecture at baseline. Following *P. chabaudi* infection, disruption of white-red pulp boundaries was observed in both genotypes; however, this disruption was more extensive in CNBP-deficient spleens, characterized by diffuse lymphoid regions and reduced architectural definition (**Fig 3A**). Furthermore, gross morphology in both infection models revealed visibly smaller spleens in CNBP-deficient mice relative to infected WT counterparts (**Fig 3B**). Although spleen size increased post-infection (pi) compared to baseline in both genotypes, splenomegaly was significantly attenuated in CNBP-deficient mice at day 9pi with *P. chabaudi* infection (**Fig 3C**). Similarly, during *P. berghei* infection, KO spleens at day 6pi were smaller than those of WT mice despite exhibiting higher peripheral parasitemia (**Fig 3D**).

**Figure 3.**
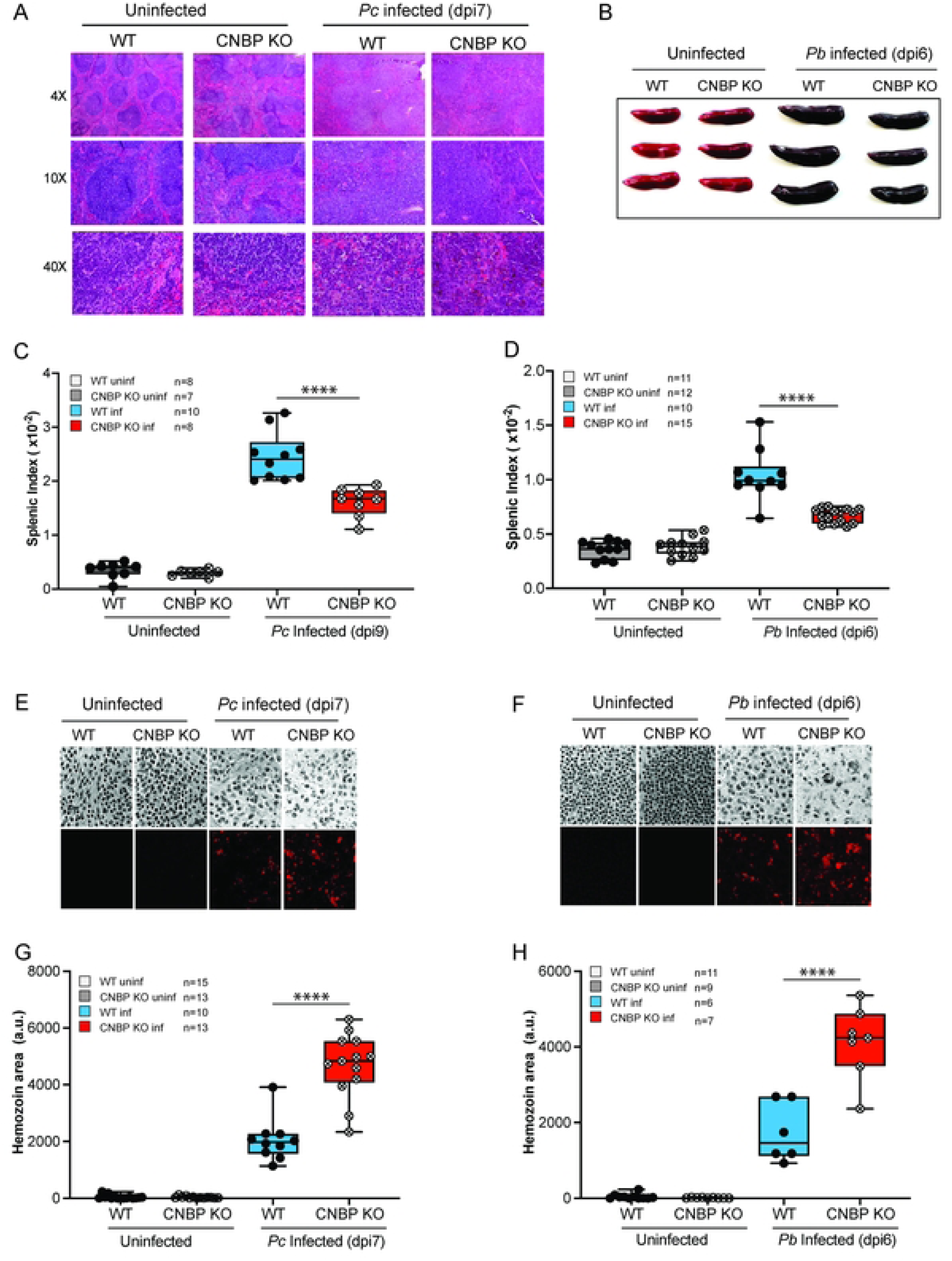
CNBP deficiency limits splenic enlargement, disrupts tissue architecture, and increases hemozoin deposition. (A) Representative HCE-stained spleen sections from *P. chabaudi*-infected WT and KO mice at 4×, 10×, and 40× magnification. WT spleens showed preserved red and white pulp organization at baseline, whereas KO spleens displayed mild architectural alterations in the uninfected state. Infection induced follicular disruption and loss of red/white pulp demarcation in both genotypes, with more pronounced disorganization in KO mice. B) Representative gross images of spleens from uninfected and *Plasmodium berghei* ANKA day 6 infected wild-type (WT) and CNBP knockout (KO) mice. Spleens were collected from both uninfected WT and KO mice, as well as from infected animals at day 6 post-infection. Images show representative mice from each group (n = 3 mice per infected group). (C-D) Splenic index (spleen weight/body weight ratio) in WT and KO mice. For *P. chabaudi*: day 0, WT (n = 8), KO (n = 7); day 9 PI, WT (n = 10), KO (n = 8). For *P. berghei* ANKA: day 0, WT (n = 11), KO (n = 12); day 6 PI, WT (n = 10), KO (n = 15). Statistical analysis was performed using two-way ANOVA with multiple-comparison testing; \*\*\*\**P* < 0.0001. Data are presented as mean ± SEM. (E-H) Quantification of splenic hemozoin deposition by reflection microscopy. For *P. chabaudi*: day 0, WT (n = 15), KO (n = 13); day 9 PI, WT (n = 10), KO (n = 13). For *P. berghei* ANKA: uninfected, WT (n = 11), KO (n = 9); day 6 PI, WT (n = 6), KO (n = 7). Hemozoin crystal area was quantified using ImageJ from ≥3 fields per section per mouse. KO spleens exhibited significantly greater hemozoin accumulation at peak infection. Statistical analysis was performed using two-way ANOVA with multiple comparisons; \*\*\*\**P* < 0.0001. Data are representative of two independent experiments (*P. chabaudi*) and one independent experiment (*P. berghei*).

Quantitative reflection confocal microscopy revealed increased hemozoin accumulation in CNBP-deficient spleens at day 9pi with *P. chabaudi* infection (**Fig 3E, 3G**) and at day 6pi with *P. berghei* infection (**Fig 3F, 3H**). Across both models, CNBP deficiency uncoupled parasite burden from splenic enlargement, with reduced splenic hypertrophy coinciding with increased pigment accumulation.

### CNBP is essential for IL12-IFN**-**γ axis activation during *Plasmodium* infection

Cytokines play a pivotal role in orchestrating the immune response to *Plasmodium* infection [**3,31]**. To determine whether CNBP regulates cytokine responses during malaria infection, we measured systemic cytokines in plasma from WT and CNBP KO mice (≥5 mice per group; both males and females) infected with *P. chabaudi* or *P. berghei*. In parallel, we assessed cytokine release from splenocytes isolated from *P. berghei* ANKA infected mice on day 6 post-infection and cultured *ex vivo*.

CNBP KO mice exhibited markedly reduced plasma IL-12β during *P. chabaudi* infection at peak parasitemia (day 7 p.i.; p < 0.0001) (**Fig 4A**). Plasma cytokines were measured longitudinally at days 0, 3, 5, 7, and 9pi to capture the temporal dynamic of cytokine response. In the *P. berghei* model, IL-12β plasma levels measured on day 6 post-infection were likewise reduced in CNBP KO mice compared with WT controls (**Fig 4B**).

**Figure 4.**
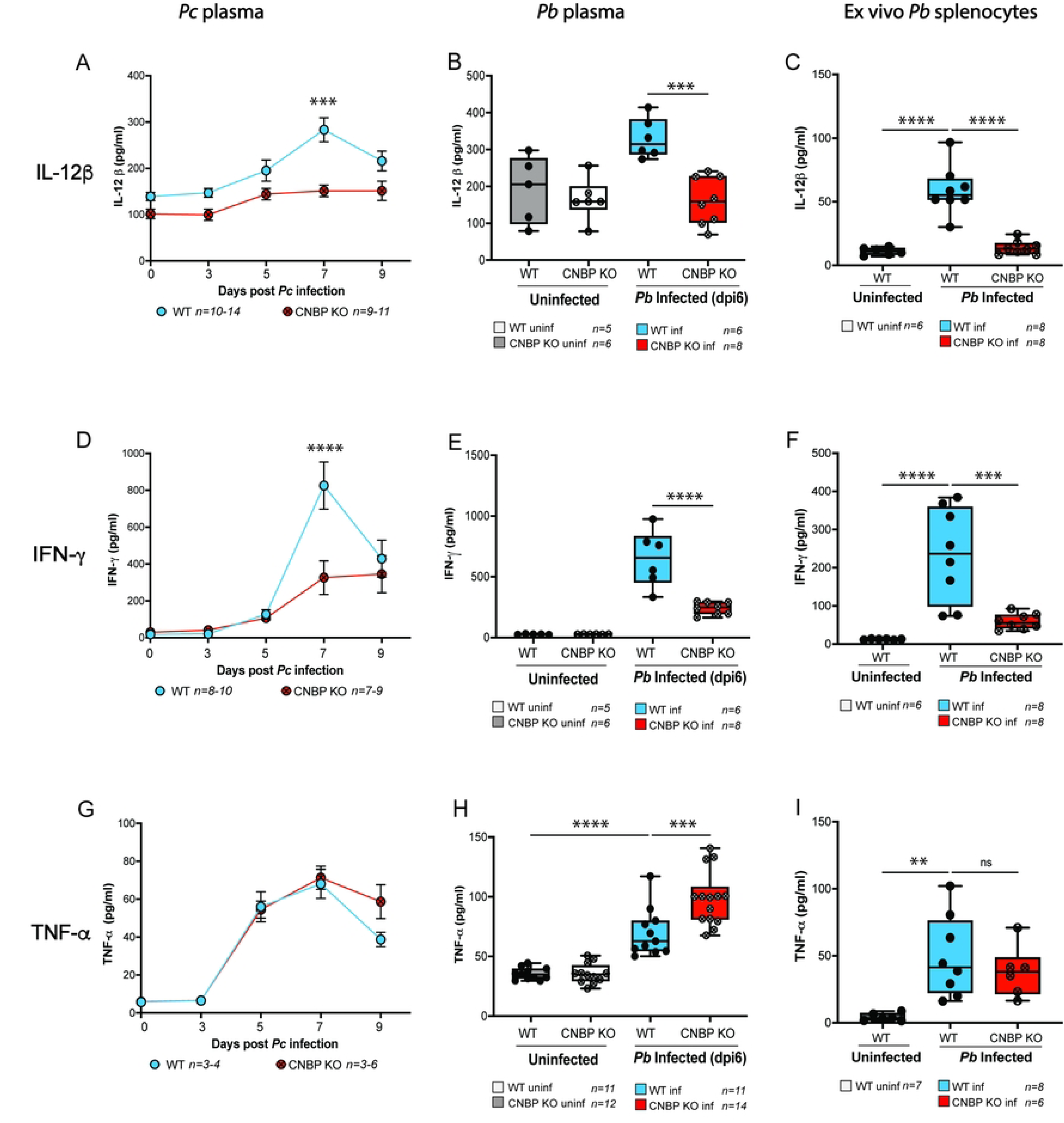
**CNBP is essential for IL12-IFN-γ axis activation during *Plasmodium* infection** (A) Plasma IL-12β levels in *P. chabaudi*-infected WT and CNBP knockout (KO) mice (n = 9-14 per group). KO mice exhibited significantly reduced IL-12β at day 7 post-infection (PI). Statistical analysis was performed using two-way ANOVA with multiple comparisons; \*\*\**P* < 0.001. (B) Plasma IL-12β levels in *P. berghei* ANKA-infected mice. Day 0: WT (n = 5-11), KO (n = 6-12); day 6 PI: WT (n = 5-17), KO (n = 7-22). KO mice showed significantly reduced IL-12β. Statistical analysis was performed using one-way ANOVA with multiple comparisons; ***P<0.001. (C) IL-12β secretion from ex vivo splenocytes cultured for 24 h. Uninfected WT (n = 6); day 5 PI WT (n = 8) and KO (n = 8). KO splenocytes secreted significantly less IL-12β compared to WT. Statistical analysis was performed using one-way ANOVA with multiple comparisons; *P* < 0.0001. (D) Plasma IFN-γ levels in *P. chabaudi*-infected WT and KO mice (n = 7-11 per group). KO mice exhibited significantly reduced IFN-γ at day 7 PI. Statistical analysis was performed using two-way ANOVA with multiple comparisons; \*\*\*\**P* < 0.0001. (E) Plasma IFN-γ levels in *P. berghei* ANKA-infected mice (same cohorts as in B). KO mice showed significantly reduced IFN-γ. Statistical analysis was performed using one-way ANOVA with multiple comparisons; \*\*\*\**P* < 0.0001. (F) IFN-γ secretion from *ex vivo* splenocytes cultured for 24 h (same cohorts as in C). KO splenocytes secreted significantly less IFN-γ than WT. Statistical analysis was performed using one-way ANOVA with multiple comparisons; ***, P<0.001; ****P<0.0001. (G) Plasma TNF-α levels in *P. chabaudi*-infected WT and KO mice (n = 3-6 per group). TNF-α levels were not significantly different between genotypes. Statistical analysis was performed using two-way ANOVA. (H) Plasma TNF-α levels in *P. berghei* ANKA-infected mice (same cohorts as in B). *P. berghei* ANKA-infected KO mice exhibited significantly elevated TNF-α levels. Statistical analysis was performed using one-way ANOVA with multiple comparisons; \*\*\**P* <0.001; \*\*\*\**P* <0.0001. (I) TNF-α secretion from *ex vivo* splenocytes cultured for 24 h (same cohorts as in C). TNF-α production was comparable between the WT and KO groups. Statistical analysis was performed using one-way ANOVA; \*\**P* < 0.01; n.s., not significant. Data is pooled from 2-3 independent experiments and are presented as mean ± SEM.

To assess whether reduced systemic IL-12β reflected altered cytokine production by immune cells, splenocytes were isolated from *P. berghei*-infected mice on day 6 post-infection and cultured *ex vivo* (1 × 10⁶ cells per well for 24 hours in RPMI). Splenocytes from uninfected mice were used as controls. Under these conditions, splenocytes from CNBP KO mice secreted substantially less IL-12β than WT splenocytes (**Fig 4C**), with a magnitude of reduction exceeding that observed in plasma. These findings indicate impaired cytokine production at the level of splenic immune cells. Together with our earlier demonstration that CNBP binds the *Il12*β promoter in MoDCs, these data support defective IL-12β production in CNBP-deficient immune cells.

Plasma IFN-γ mirrored the IL-12β phenotype, with significantly reduced concentration in CNBP KO mice during both *P. chabaudi* and *P. berghei* infections (**Fig 4D, 4E**). Consistently, *ex vivo* splenocyte cultures from CNBP KO mice secreted lower amounts of IFN-γ than WT (**Fig 4F**), indicating impaired activation of the IL12-IFN-γ axis in the absence of CNBP.

TNF-α levels displayed model-specific dynamics. During *P. chabaudi* infection, plasma TNF-α levels were comparable between WT and CNBP KO mice (**Fig 4G**). In contrast, during *P. berghei* infection, plasma TNF-α levels were elevated in CNBP KO mice at 6dpi (**Fig 4H**), However, TNF-α secretion from *ex vivo* splenocyte cultures did not differ between genotypes (**Fig 4I**).

Systemic IL-6 levels were similar between WT and KO mice in both infection models (**Fig S4A, S4B**). By contrast, *ex vivo* splenocyte cultures from CNBP KO mice exhibited reduced IL-6 secretion relative to WT controls (**Fig. S4C**), indicating impaired cytokine production by immune cells that is not reflected at the systemic level. Plasma IL-10 levels were significantly elevated in CNBP KO mice during *P. chabaudi* infection at day 9 p.i. (**Fig S4D**), whereas IL-10 levels during *P. berghei* infection and in *ex vivo* splenocyte cultures were comparable between genotypes (**Fig S4E, S4F**). Plasma IL-18 levels and *ex vivo* IL-18 secretion were unchanged in CNBP KO mice in both infection models (**Fig S4G, S4H, S4I**), indicating that CNBP does not broadly regulate inflammasome-associated cytokine production under these conditions.

Collectively, our results show that CNBP deficiency selectively impaired IL-12β and IFN-γ responses across both malaria models, highlighting a critical role for CNBP in promoting the IL12β- IFN-γ axis. Model-specific alterations, including TNF-α elevation in *P. berghei* at 6dpi and late IL-10 induction (dpi9) in *P. chabaudi*, likely reflect differential innate immune activation between chronic non-lethal (*P. chabaudi*) and acute cerebral (*P. berghei*) malaria. These cytokine defects are consistent with the altered parasitemia and survival outcomes and indicate that CNBP contributes to shaping both the magnitude and context of inflammatory immune responses during malaria infection.

### CNBP deficiency is associated with selective alterations in splenic gene expression and cellular remodeling in the spleen during *Plasmodium chabaudi* infection

To obtain an unbiased view of CNBP-dependent immune regulation, independent cohorts of WT and CNBP-deficient mice (8-10 weeks old) were infected intraperitoneally with 1× 10^6^ *P. chabaudi*-AS infected erythrocytes. Spleens were harvested at days 0, 3, 5, 7, and 9 post-infection for bulk RNA sequencing.

Global transcriptional responses to infection were largely similar between genotypes. CNBP transcripts were undetectable in knockout mice, confirming efficient deletion in hematopoietic cells. In WT mice, CNBP expression was dynamically regulated during infection, increasing from baseline and peaking sharply at day 7 p.i., coincident with maximal parasitemia, consistent with CNBP induction as part of the host stress-responsive immune program (**Fig 5A**).

**Figure 5.**
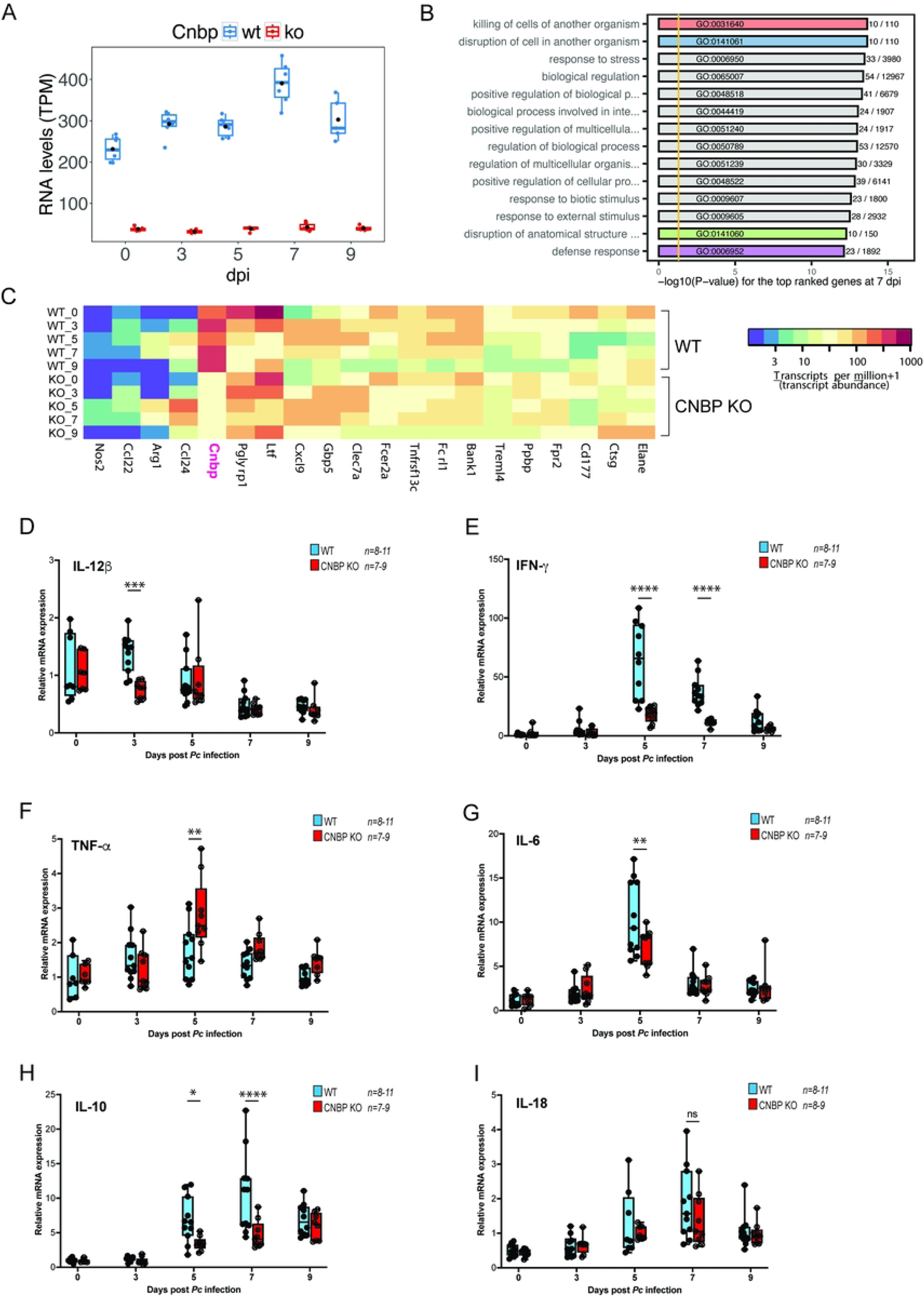
Cnbp deficiency alters RNA expression of cytotoxic genes during *P. chabaudi* infection. (A) RNA Expression levels (transcripts per million) for Cnbp at 0 to 9 days post infection (dpi). Expression levels for individual mice are represented as blue (wild type) and red (*Cnbp* KO) circles. The means are represented as black circles. (B) GO enrichment analysis for the top-ranked genes at 7 dpi. Genes were ranked by absolute fold-change between wild type and KO at 7 dpi, and all genes with a 2-fold change or greater (n = 89) were tested for enriched GO terms in the biological process category. Four of the significantly enriched GO terms (shown in red, blue, green, and purple) have a potential role in cell cytotoxicity and the defense response. The fully expanded term for GO:0141060 is ‘disruption of anatomical structure in another organism’. (C) RNA expression levels among 20 significant immune genes. The genes in the heatmap (bottom) have a 2-fold or greater differential expression between wild type and KO at one or more time points. The 10 genes belonging to GO:0031640, GO:0141061, and GO:0141060 (panel B) are *Nos2*, *Ccl22*, *Arg1*, *Ltf*, *Gbp5*, *Clec7a*, *Fcer2a*, *Ppbp*, *Ctsg*, and *Elane*. *Pglyrp1* and *CxclS* have a 2-fold differential expression at 9 dpi. The *Cnbp* gene (labeled Pink) is included for comparison. The remaining 7 genes are also immune genes. (D-I) RT-qPCR analysis was performed on total RNA isolated from spleens of WT and CNBP KO mice collected at days 0, 3, 5, 7, and 9 post-infection (p.i.). Relative expression of *Il12b, Ifn-*γ*, Tnf-*α*, Il-c, Il-10,* and *Il18* was determined. *Il12b* transcripts were significantly reduced in CNBP-deficient mice at day 3 p.i.; Ifn-γ at days 5 and 7 p.i.; Il6 at day 5 p.i.; and Il10 at day 5 and 7 p.i. In contrast, Tnf-α was upregulated in CNBP KO mice, with Il18 unchanged, if not mildly elevated. - between genotypes at any time point. Gene expression levels were normalized to Hprt and calculated using the 2^–ΔΔCt^ method. Data represent mean ± SEM (n > 5 mice per group per time point). Statistical comparisons between genotypes at each time point were performed using an unpaired two-tailed Student’s t-test. *P*-values are indicated as follows: \**P* < 0.05, \*\**P* < 0.01, \*\*\**P* < 0.001, \*\*\*\**P* < 0.0001.

Gene ontology (GO) enrichment analysis on the top-ranked genes (≥2-fold differential expression) revealed significant enrichment for cytotoxicity-related processes (GO:0031640, GO:0141061, and GO:0141060) (**Fig 5B**). Ten genes with known cytotoxic functions (Nos2, Ccl22, Arg1, Ltf, Gbp5, Clec7a, Fcer2a, Ppbp, Ctsg, and Elane) were differentially expressed at day 7, while two additional cytotoxic genes (Pglyrp1 and Cxcl9) differentially expressed at day 9 (**Fig 5C**). Il1b transcripts were elevated at day 3, Gzma at day 5, and Ccl24 just prior to peak parasitemia at day 5 (**Fig S5A**). These data depict that CNBP-dependent transcriptional changes are enriched in cytotoxic and defense-associated transcriptional programs. These alterations reflect a compensatory response to counterbalance impaired IL12-IFN-γ signaling.

To determine whether CNBP-dependent transcriptional changes were accompanied by alterations in immune cell composition, we quantified splenic dendritic cells and T cell subsets throughout the course of *P. chabaudi* infection (days 0, 3, 5, 7, and 9 post-infection). CD11c⁺ MHC-II⁺ dendritic cells were assessed within the CD45⁺ compartment by flow cytometry. Frequencies of dendritic cells remained comparable between WT and CNBP-deficient mice at all time points (**Fig S5A**), indicating that CNBP deficiency does not measurably affect splenic dendritic cell abundance during infection. In contrast, analysis of T cell subsets revealed genotype-dependent differences emerging at the late stage of infection. At day 9 post-infection, CNBP-deficient mice exhibited a significant reduction in CD4⁺ T cells (CD45⁺ CD3⁺ CD4⁺) accompanied by a relative increase in CD8⁺ T cells (CD45⁺ CD3⁺ CD8⁺) compared with WT controls (**Fig S5B, S5C**). No differences in T cell frequencies were observed at earlier time points, suggesting that these alterations arise because of ongoing infection rather than reflecting baseline developmental defects.

Although RNA-seq captured global transcriptional remodeling, canonical malaria-protective cytokines, including Il12β, Ifn-γ, Il6, Il10, Tnf-α, and Il18 were expressed at low levels in whole spleen RNA and were not reliably identified as differentially expressed (**Fig S5D**). To address this, we performed targeted qPCR in independent cohorts of WT and CNBP-deficient mice at days 0, 3, 5, 7, and 9 post-infection (n = 7-11 per group).

IL-12β transcripts were significantly reduced in CNBP-deficient spleens as early as day 3 p.i., with Ifn-γ also diminished at days 5-7 (**Fig 5D, 5E**), consistent with reduced activity of the canonical IL-12-IFN-γ axis. Il6 and Il10 transcripts were selectively reduced at day 5 and day 7, respectively (**Fig 5G, 5H**), while Tnf-α and Il18 expression remained comparable across genotypes (**Fig 5F, 5I**) suggesting that CNBP deficiency is associated with selective effects on a subset of cytokine pathways.

## Discussion

Effective immunity to blood-stage malaria requires tight coordination between innate sensing, transcriptional control of cytokine programs, and adaptive immune remodeling **[32,33]**. While pattern recognition pathways initiating these responses have been extensively studied, how parasite-derived cues are integrated at the level of nuclear transcriptional regulators remains incompletely understood. Genetic and immunological studies in both murine models and humans have consistently demonstrated that the IL12-IFN-γ axis is central to Th1 polarization, immune cell activation, and splenic clearance of infected erythrocytes, yet the transcriptional mechanisms selectively sustaining this pathway without provoking immunopathology remain poorly defined **[2,3]**. In this study, we identify Cellular Nucleic Acid-Binding Protein (CNBP) as a conserved, stimulus-responsive transcriptional regulator that couples *Plasmodium* sensing to IL12-IFN-γ axis activation, splenic immune architecture, resulting in parasite control *in vivo* (Fig 6)

**Figure 6.**
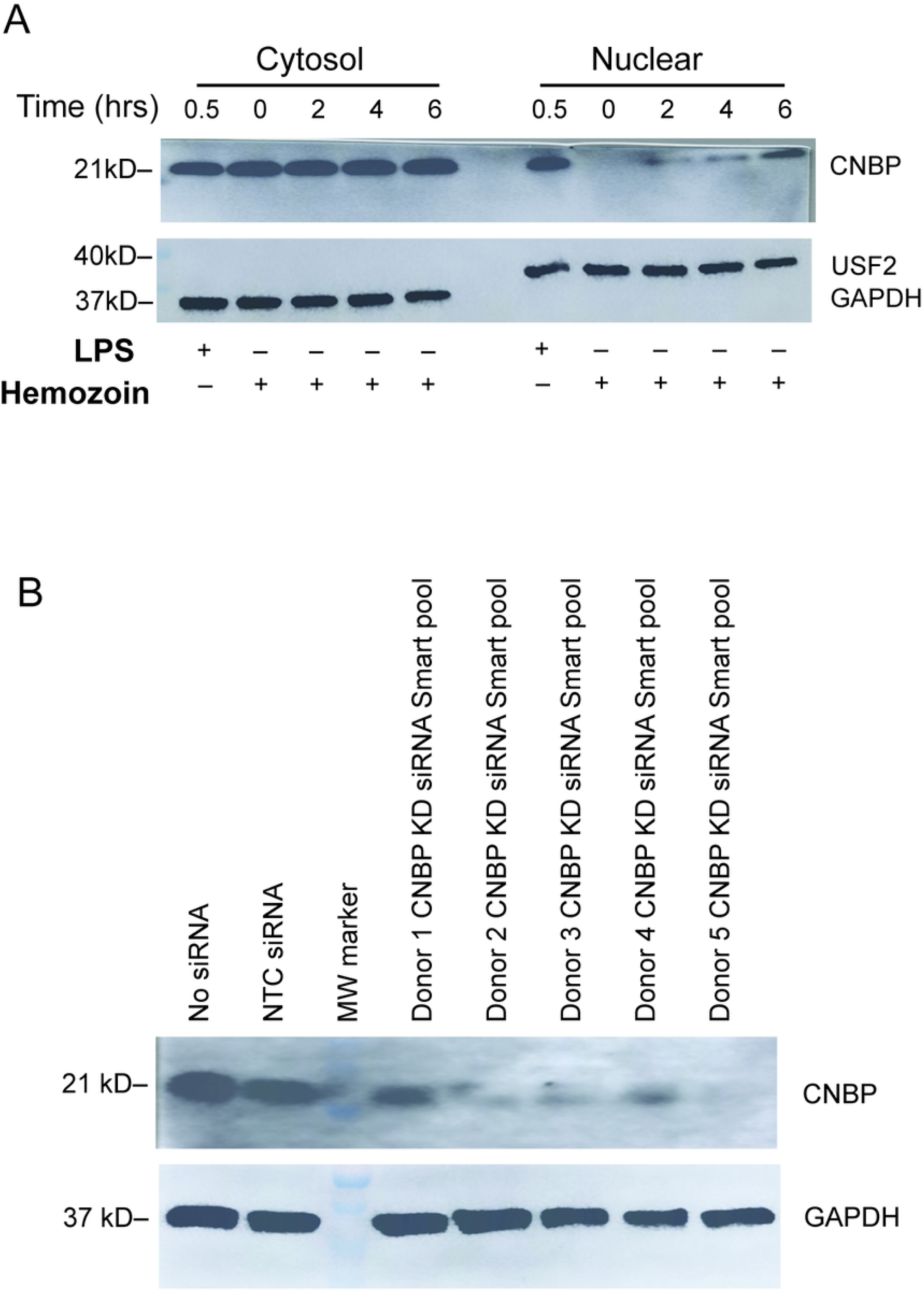
CNBP orchestrates protective IL-12-IFN-γ responses in malaria via dendritic cell activation and T cell polarization. Schematic model illustrating CNBP function during blood-stage malaria. Upon exposure to *Plasmodium falciparum*-infected red blood cells (iRBCs), CNBP translocates from the cytosol of DCs into the nucleus, where it promotes transcriptional activation of Il12b (encoding IL-12β). This drives IL-12 production and IFN-γ secretion by CD4⁺ T cells, supporting a robust Th1 proinflammatory response essential for parasite control and clearance. Loss of CNBP leads to markedly reduced IL-12 and IFN-γ levels, impaired Th1 polarization, and a skewed splenic T cell composition favoring CD8⁺ over CD4⁺ subsets. Consequently, parasitemia increases, hemozoin accumulation intensifies, splenic microarchitecture becomes disrupted, and infection recrudesces. This model highlights CNBP as a critical innate transcriptional amplifier that links innate sensing of *Plasmodium* to effective adaptive immunity and tissue homeostasis.

Our data establish that CNBP undergoes rapid and stimulus-dependent nuclear translocation in both murine and human dendritic cells following exposure to *Plasmodium* infected erythrocytes. This response is selective, as uninfected erythrocytes fail to induce translocation, and is recapitulated by hemozoin, a dominant malaria associated danger signal known to engage innate immune sensors **[12,13,34]**. These findings extend prior work linking CNBP to bacterial and viral immunity by demonstrating that CNBP is not pathogen restricted but instead functions as a broader transcriptional integrator of microbial and parasitic signals. CNBP is regulated by phosphorylation downstream of TLRs. The kinase TAK1 is important in controlling phosphorylation [**22]**. These events in turn trigger nuclear accumulation of CNBP.

Mechanistically, here we show that the nuclear accumulation of CNBP correlates with direct engagement of the *Il12*β promoter at defined upstream regulatory regions following parasite stimulation. The inducible nature of this binding, together with the loss of IL-12β transcription and secretion in CNBP-deficient dendritic cells, supports a model in which CNBP acts as a transcriptional amplifier rather than a basal activator of *Il12*β. This is consistent with earlier reports showing that CNBP cooperates with inducible transcription factors downstream of innate signaling rather than functioning as a standalone DNA binding factor **[24]**. Importantly, the conservation of this regulatory axis in human MoDCs underscores its translational relevance.

*In vivo*, CNBP deficiency resulted in impaired control of blood stage parasitemia across both non-lethal and lethal malaria models. The recrudescence observed during *P. chabaudi* infection is particularly informative, as it resembles phenotypes reported in models of defective Th1 priming or premature effector contraction **[31]**. Rather than reflecting an inability to initiate immune responses, these kinetics suggest a failure to maintain protective immunity during peak and post-peak infection phases. This interpretation is consistent with the early reduction in *Il12*β expression, followed by diminished IFN-γ production and late-stage CD4⁺ T cell loss.

Importantly, CNBP deficiency did not result in broad suppression of inflammatory pathways. Inflammasome-associated cytokines such as IL-18 were unaffected, and model-specific increases in TNF-α or IL-10 likely reflect compensatory or regulatory responses rather than direct CNBP control. This pathway selectivity mirrors prior observations that partial disruption of the IL12-IFN-γ axis can uncouple parasite burden from inflammatory pathology, particularly in cerebral malaria models, where excessive Th1 responses exacerbate tissue damage [**35,36]**. Together, these data position CNBP as a pathway selective regulator essential for sustaining Th1-biased immunity rather than as a global amplifier of inflammation.

A striking feature of CNBP deficient mice was the dissociation between elevated parasitemia and reduced splenomegaly. Splenic enlargement during malaria is generally considered an adaptive response, reflecting myeloid expansion, enhanced filtration capacity, and immune architectural remodeling necessary for efficient clearance of infected erythrocytes **[37]**. The attenuated splenomegaly observed in CNBP-deficient mice, therefore, indicates a failure of functional immune expansion rather than protection from inflammation.

This impaired remodeling coincided with a surprisingly increased hemozoin accumulation and greater architectural disarray, suggesting defective processing of parasite debris. Given the well-established role of IFN-γ in licensing macrophage phagocytic and degradative programs [**38]**, reduced IFN-γ signaling in CNBP-deficient mice likely compromises hemozoin clearance, reinforcing pigment accumulation and disrupting splenic function. The observation that hemozoin itself induces CNBP nuclear translocation further supports a feed-forward regulatory loop in which accumulating parasite products normally reinforce IL-12 dependent immune organization, a loop that is broken in the absence of CNBP.

Systemic and *ex vivo* cytokine analyses demonstrate that CNBP deficiency selectively impairs IL-12β and IFN-γ production across both malaria models, while leaving inflammasome-associated cytokines, such as IL-18, largely intact. While IFN-γ production has been shown to be potentially lethal in P chabaudi infection, IL-12 deficiency has a phenotype considerably closer to that which we observed in CNBP deficient mice. One potential explanation is that the regulation of IFN-γ production in NK cells might be independent of CNBP unlike that from lymphocytes, although NK cels from IL12 knockouts appear to be severely restricted in their ability to produce IFN-γ **[39]**. This specificity argues against a generalized defect in innate activation and instead positions CNBP as a pathway selective regulator.

Transcriptomic analysis revealed that, in the absence of CNBP, the immune system adopts an alternative transcriptional strategy characterized by enrichment of cytotoxic and antimicrobial gene programs. Although bulk RNA-seq lacks sensitivity for low-abundance cytokine transcripts, a limitation addressed by targeted qPCR **[40]**, its strength lies in capturing broader immune reprogramming. The skewing toward cytotoxicity associated genes, together with increased CD8⁺ T cell representation and granzyme expression, suggests compensatory engagement of non-canonical effector pathways when Th1 immunity is compromised. The upregulation of genes associated with neutrophil activity, myeloid cytotoxicity, and oxidative stress suggests a shift toward innate defense mechanisms that may partially restrain infection but are insufficient to restore effective parasite clearance.

Despite its central role in dendritic cell cytokine output, CNBP deficiency did not alter splenic dendritic cell abundance during infection, indicating that CNBP primarily regulates function rather than development or survival of these cells. In contrast, CNBP-deficient mice exhibited late-stage alterations in T cell subset distribution, characterized by reduced CD4⁺ T cells and a relative increase in CD8⁺ T cells. Importantly, these cellular changes emerge after the onset of cytokine defects, supporting a model in which altered immune architecture is a downstream consequence of impaired CNBP-dependent transcriptional programming rather than its primary cause.

One of the more intriguing findings of this study is the modest prolongation of survival observed in CNBP-deficient mice during experimental cerebral malaria, despite higher parasitemia. This apparent paradox aligns with established paradigms in which partial attenuation of IL12-IFN-γ or TNF-α signaling can reduce lethal immunopathology at the cost of impaired parasite clearance **[36,41]**. CNBP deficiency therefore appears to shift the host-pathogen equilibrium toward a more tolerable inflammatory state, highlighting its role as a molecular node that balances protective immunity against inflammatory damage. While our data do not directly assess cerebral pathology, these findings strongly suggest that CNBP influences not only resistance but also disease tolerance mechanisms during malaria.

Although our *in vitro* human data are currently limited to observations of CNBP nuclear translocation and transcriptional activation, they nonetheless provide an important mechanistic anchor linking parasite recognition to cytokine gene regulation. The immediate upstream signaling events that trigger CNBP mobilization in human dendritic cells remain undefined and warrant further investigation. In addition, while our data support a central role for CNBP in dendritic cell-driven IL12-IFN-γ programming, we cannot fully exclude contributions from other immune compartments, including monocytes or NK cells. Because our knockout strategy targets the hematopoietic compartment broadly, future studies using cell type-specific conditional deletions will be required to more precisely define cell-intrinsic functions of CNBP. Finally, in the *P. berghei* model, correlating survival outcomes with brain histopathology and local cytokine profiling will be important to clarify whether CNBP deficiency alters cerebral malaria pathology.

Collectively, our results establish CNBP as a stimulus-responsive transcriptional regulator that couples *Plasmodium* sensing to IL-12-IFN-γ axis activation, splenic immune organization, and disease outcome. By sustaining Th1-polarizing cytokine programs and immune architecture without altering dendritic cell abundance, CNBP enables effective parasite control and coordinated inflammatory responses. Its absence leads to selective transcriptional and cytokine defects, late-stage immune reprogramming toward compensatory cytotoxic pathways, impaired splenic remodeling, and persistent parasitemia, revealing CNBP as a critical determinant of how host immunity balances resistance and tolerance during malaria.

These findings expand the conceptual framework of malaria immunity by highlighting transcriptional co-regulators, not only pattern-recognition receptors, as critical determinants of host defense and disease trajectory. More broadly, they position CNBP as a potential target for modulating inflammatory balance in malaria and other infectious diseases where IL12-IFN-γ signaling plays a central role.

## Materials and methods

### Ethics Statement

All animal experiments were conducted in accordance with the NIH Guide for the Care and Use of Laboratory Animals and the standards of the American Association for Laboratory Animal Science (AALAS). Protocols were approved by the Institutional Animal Care and Use Committee (IACUC) at the University of Massachusetts Chan Medical School (PROTO202000093).

Human leukocytes were obtained as de-identified leukopacks through the UMass Chan Leukocyte Research Core. Samples were originally collected by the Rhode Island Blood Center, a non-profit blood center that is a division of the New York Blood Center.

### Mice

C57BL/6J mice were obtained from The Jackson Laboratory (Strain #:000664). CNBP^fl/fl^ mice crossed to *Vav-iCre* transgenic mice were generated and maintained in house. Mice were housed under specific pathogen free conditions at the UMass Chan Medical School animal facility, maintained on a 12h light/dark cycle, and provided autoclaved chow and sterile water ad libitum. Experimental cohorts consisted of male and female mice aged 8-10 weeks. Unless otherwise indicated, data was pooled from both sexes.

### Genotyping

Genotyping was performed by Transnetyx (Cordova, TN) using standard real-time PCR-based assays.

### Plasmodium Parasites and Experimental Infection

GFP expressing *Plasmodium chabaudi AS* and *Plasmodium berghei ANKA* parasites were expanded in C57BL/6 mice prior to experimental use. Frozen parasite stocks were passaged at least twice to ensure stable parasitemia before experimental infection. Mice were infected intraperitoneally with 1 × 10⁶ infected red blood cells (iRBCs) diluted in sterile PBS. Control animals received sterile PBS alone.

### Parasitemia, Blood Glucose, and Survival Monitoring

#### Parasitemia

Parasitemia was quantified either by Giemsa-stained thin blood smears and by flow cytometry using GFP-expressing parasites. Giemsa staining was used for morphological confirmation, while flow cytometry enabled higher throughput quantitative assessment.

#### Blood Glucose

Peripheral blood glucose levels were measured on alternate days using a handheld glucometer (& NovaMax strip) with tail vein blood collected at consistent times to minimize circadian variation.

#### Survival Analysis

For survival studies*, P. berghei ANKA* infected mice were monitored daily. Animals reaching predefined humane endpoints or experiencing >20% body weight loss were euthanized in accordance with IACUC guidelines. Survival studies included n= 25-30 mice per group. Survival curves were analyzed using the Mantel Cox log-rank test.

### Flow Cytometry

#### Peripheral Blood

Tail vein blood (2–3 µL) was collected into ice-cold HBSS supplemented with 2% FBS and fixed with 0.025% glutaraldehyde. Samples were stained with anti-Ter119 PE-Cy7 (Invitrogen, eBioscience; Cat no.25592182, Dilution 1:50). GFP⁺ events within the Ter119⁺ gate was quantified **[42]**. Gating strategy is shown in Fig S2d.

#### Splenocytes

Single cell splenocyte suspensions were prepared by mechanical dissociation through a 70 µm strainer, followed by red blood cell lysis using ACK buffer. Cells were stained with viability dye (Ghost 510; Tonbo Biosciences) and fluorophore conjugated antibodies to CD45, CD3, CD4, CD8α, CD11c, and MHC class II (I-A/I-E). In a selected experiment, CD19, CD11b, and TCRβ were also included (Fluorophore and dilution information in Sup. Table S3). Doublets were excluded by FSC-A/FSC-H gating. Data were acquired on a Bio-Rad ZE5 Cell Analyzer and analyzed using FlowJo software (Version 10.10.0).

#### Splenic Index and Histology

Spleens were harvested, blotted dry, and weighed. The splenic index was calculated as:

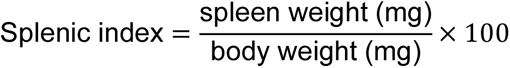

For histological analysis, spleens were fixed in formalin, paraffin embedded, sectioned, and stained with hematoxylin and eosin. Hemozoin deposition was visualized using reflection microscopy and quantified using ImageJ software with consistent thresholding parameters across samples.

### Cell Isolation and Culture

#### Murine Splenocytes and BMDCs

Splenocytes were cultured in RPMI-1640 supplemented with 10% FBS and 1% penicillin/streptomycin. Bone marrow–derived dendritic cells (BMDCs) were generated from femoral and tibial marrow and differentiated in the presence of rM-CSF (Bio legend Cat no. 576406, 50ng/ml) for 5-7 days, with partial medium replacement every 2-3 days.

#### Human PBMCs and Monocyte-Derived DCs

PBMCs were isolated by density gradient centrifugation. CD14⁺ monocytes were purified by negative selection (Miltenyi PAN Monocyte Isolation Kit, Cat. no.130-096-537) and differentiated into dendritic cells in RPMI-1640 containing rGM-CSF (Bio legend Cat no. 572903, 50ng/ml) and IL-4 (Bio legend Cat no. 574004; 50ng/ml) for 7 days. Cells were derived from five independent healthy donors.

### Cell Stimulation and Fractionation

Murine or human dendritic cells were stimulated with *Plasmodium falciparum* iRBCs (from ATCC, Cat no. mra-102; ratio 1:10), purified natural hemozoin (100 µM), or ultrapure LPS (100 ng/mL; InvivoGen, cat. no. tlrl-eblps). Hemozoin preparations were done according to Parroche P. et al 2006 **[14].**

Nuclear and cytoplasmic extracts were prepared using the NE-PER Nuclear and Cytoplasmic Extraction Kit (Thermo Fisher Scientific, Cat. no. 78835) according to the manufacturer’s instructions.

### Immunoblotting and Immunofluorescence

#### Immunoblotting

Equal amounts of cytosolic or nuclear protein (15 µg) were resolved by SDS-PAGE and transferred to PVDF membranes. Membranes were probed with antibodies against CNBP H-7 (Santa Cruz, Cat. no.SC-515387-X, Dilution 1:1000), USF2 (Novus Biologicals, Cat. no. NBP1-92649, Dilution 1:2000), and GAPDH D16H11-XP (CST, Cat. no. 5174S, dilution 1:2500). Signals were detected using HRP-conjugated secondary antibodies (CST, Cat no. 7076S, 7074S) and chemiluminescence.

#### Immunofluorescence

Cells were fixed with 4% paraformaldehyde, permeabilized with 0.1% Triton X-100 in PBS, blocked with 5% BSA, and stained with fluorophore-conjugated anti-CNBP antibody (CNBP-AF647, Santa Cruz, Cat no.sc-515387, dilution 1:100). Nuclei were counterstained with DAPI and imaged using a Leica SP2 confocal microscope.

### RNA Isolation, qRT-PCR, and RNA Sequencing

Total RNA was extracted Qiagen RNasey Kit (Cat. no. 74106) with on-column DNase digestion. RNA concentration was checked by Nanodrop, and integrity was assessed using a Fragment Analyzer (UMass Molecular Biology Core). cDNA synthesis was performed using *iScript* Reverse Transcription Supermix (Bio-Rad). qRT-PCR was performed using Fast SYBR Green Master mix (Applied Biosystem Cat. no. 4385617) on a Quant Studio 3 system. Gene expression was normalized to HPRT and calculated using the 2^-ΔΔCt method. Primer sequences are listed in Sup. Table S1.

For RNA sequencing, libraries were prepared using the Illumina mRNA Library Prep Kit and sequenced on a NovaSeq X Plus platform **[43]**. RNA-seq was performed on splenic RNA from n = 5 individual mice per genotype per time point.

#### Analysis of RNA-Seq data

We used Bowtie2 **[44]** to align RNA-Seq reads to the mouse genome (assembly GRCm38 / mm10, Ensemble version 100). The RSEM package **[45]** was used to calculate transcripts per million (TPM) for reads mapped to each sample. We used the EBSeq-HMM Package **[46]** to identify genes that were differentially expressed over each time course experiment. Here, we controlled for false discovery at a rate of 5%, which corresponds to a posterior probability less than 0.05 for a constant path of expression along the time course. We also calculated fold change values between the two genetic backgrounds (wildtype and Cnbp KO) at each timepoint. These fold change values were used to rank order all genes with posterior probabilities less than 0.05 for a constant expression path in one- or both-time courses. For GO enrichment analysis **[47]**, the top ranked genes (N=89, all genes were a minimum of 2-fold) were analyzed. Custom plots were generated within the R software environment **[48]** and the ggplot2 package **[49].** The RNA-Seq data and associated protocols are available from the Array-Express public repository (E-MTAB-15402).

#### Chromatin Immunoprecipitation

ChIP was performed in BMDCs stimulated with iRBCs for 2h or LPS (100 ng/ml, 30 min). Cells were crosslinked with 1% formaldehyde in PBS, and chromatin was sonicated to 100-500 bp fragments. Immunoprecipitation was performed using anti-CNBP antibody or isotype matched IgG control as shown in **[24]**

Enrichment at the *Il12b* promoter was assessed by qPCR using primers spanning -100 to -500 bp relative to the transcription start site. ChIP signals were normalized to input DNA and expressed as percent input. ChIP experiments were performed from three independent donors - each donor sample subdivided into mock, LPS, *iRBC*, and IgG control conditions, with qPCR performed in duplicates.

#### siRNA-Mediated Knockdown of CNBP

Human monocyte-derived DCs were Lipofectamine transfected with pooled CNBP specific SMART pool siRNA or non-targeting control siRNA (Dharmacon, Cat no. L-019778-00-0010) according to the manufacturer’s instructions. Knockdown efficiency (approximately 70-85%) was confirmed by immunoblotting prior to downstream analyses.

#### ELISA

Cytokine concentrations in plasma and culture supernatants were quantified using commercial ELISA kits for mouse or human IL-6, IL-10, TNF-α, IFN-γ, IL-18, and IL-12β (Sup. Table S2). Samples were analyzed in duplicate according to manufacturers’ instructions.

#### Statistical Analysis

Statistical analyses were performed using GraphPad Prism (Version 10.4.1). Two-group comparisons were performed using unpaired two-tailed t-tests. Multi-group comparisons were performed using one- or two-way ANOVA with appropriate post-hoc testing. Survival curves were analyzed using the Mantel Cox log-rank test. A P-value <0.05 was considered statistically significant.

## Acknowledgments

We thank Zhaozhao Jiang for assistance with reagent access, protocol troubleshooting, and technical support during laboratory procedures. We acknowledge Kristy Chiang for helpful input on flow cytometry panel design. We also thank Melanie Trombly for proofreading and editorial assistance on the manuscript.

## Author contributions

Conceptualization: KAF, DTG, RR; Methodology: RR, LS; Investigation: RR, LS, SB; Formal analysis: RR, LS, EKJ, KAF, DTG; Data curation: RR; Bioinformatic analysis: DC; Writing original draft: RR, KAF, DTG; Reviewing/editing: RR, EKJ, SB, KAF, DTG; Funding, Resources & Supervision: KAF, DTG.

## Data Availability

All data generated or analyzed during this study are included in this published article and its supplementary information files. RNA-Seq data have been deposited in the Array-Express public repository (E-MTAB-15402).

---

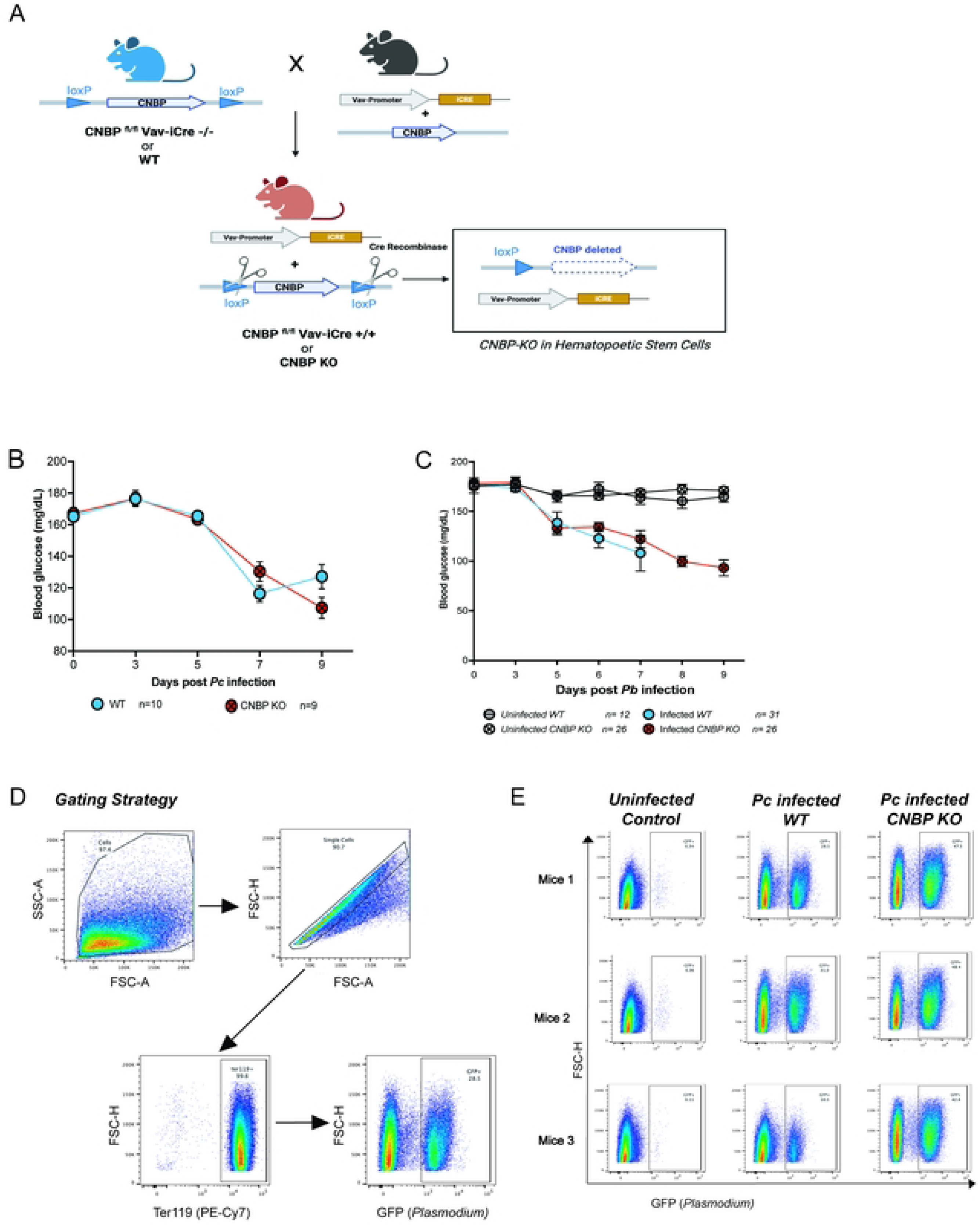

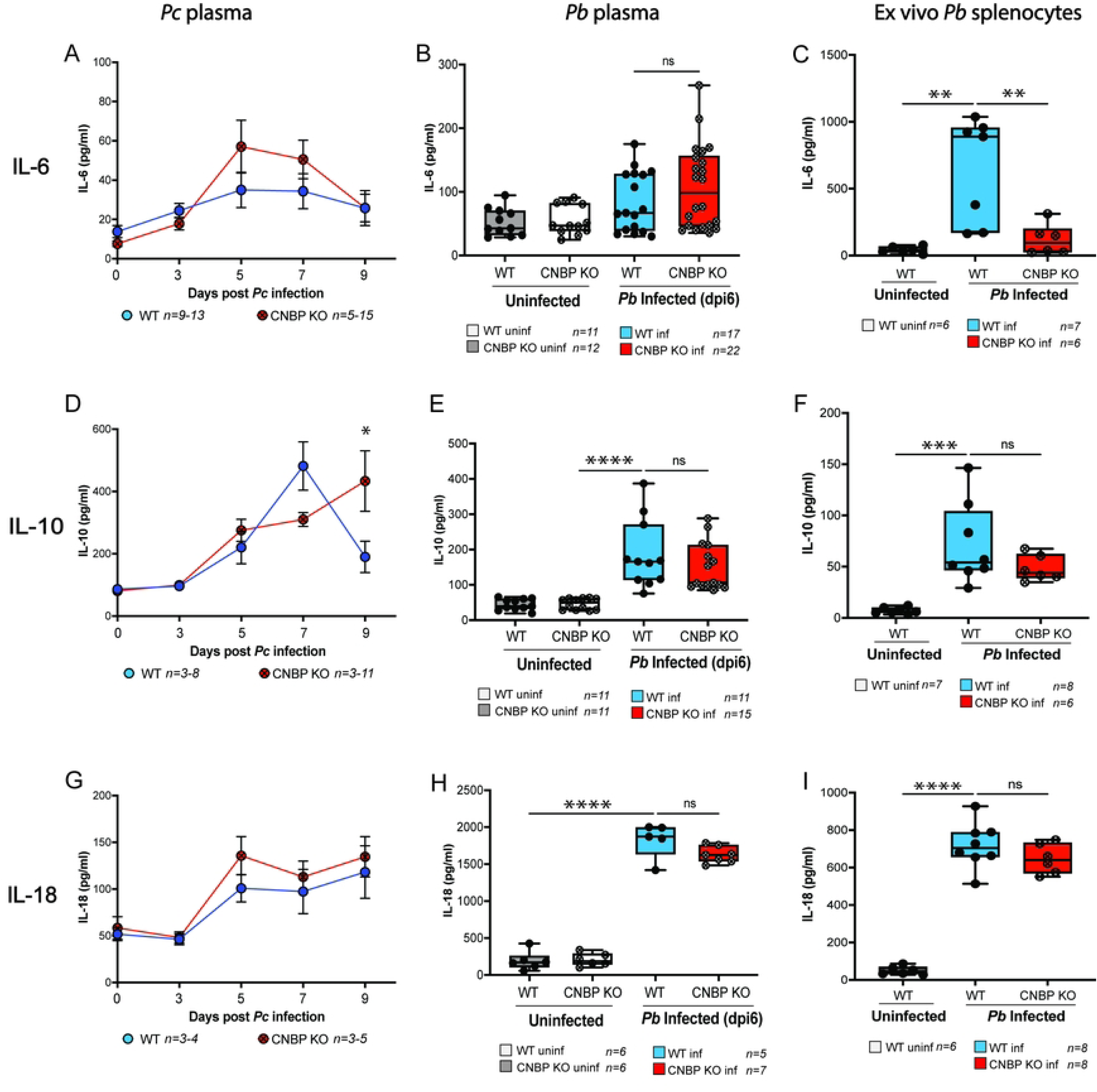

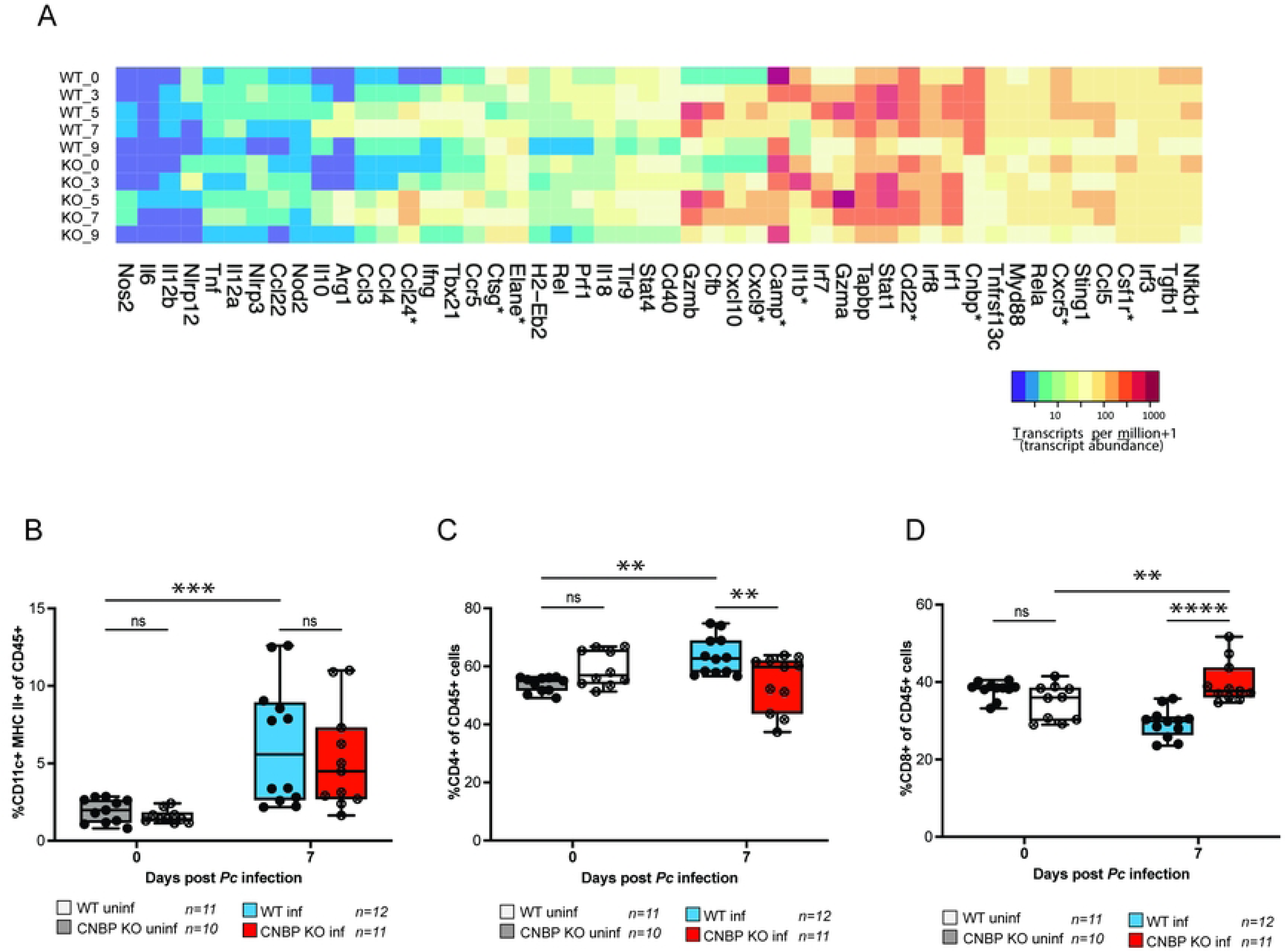

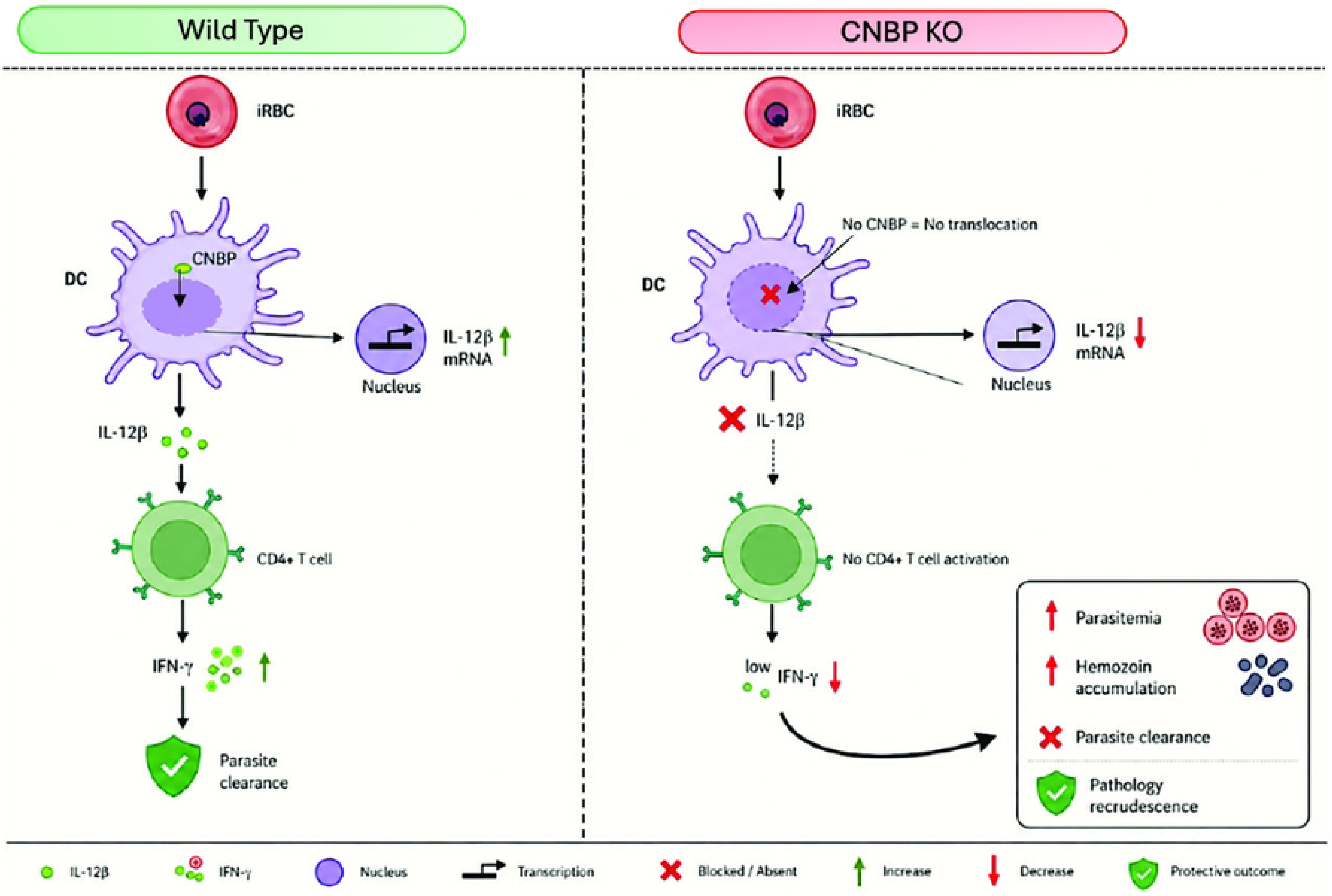

## References

1. World Health Organization. World Malaria Report (2025). Available online at: https://www.who.int/teams/global-malaria-programme/reports/world-malaria-report-2025

2. Gowda DC, Wu X. Parasite Recognition and Signaling Mechanisms in Innate Immune Responses to Malaria. Front Immunol. 2018 Dec 19; 9:3006. doi: 10.3389/fimmu.2018.03006. PMID: 30619355; PMCID: PMC6305727.

3. Langhorne J, Ndungu FM, Sponaas AM, Marsh K. Immunity to malaria: more questions than answers. Nat Immunol. 2008 Jul;9(7):725–32. doi: 10.1038/ni.f.205. PMID: 18563083.

4. Seder RA, Gazzinelli R, Sher A, Paul WE. Interleukin 12 acts directly on CD4+ T cells to enhance priming for interferon gamma production and diminishes interleukin 4 inhibition of such priming. Proc Natl Acad Sci U S A. 1993 Nov 1;90(21):10188–92. doi: 10.1073/pnas.90.21.10188. PMID: 7901851; PMCID: PMC47739.

5. Stevenson MM, Urban BC. Antigen presentation and dendritic cell biology in malaria. Parasite Immunol. 2006 Jan-Feb;28(1-2):5-14. doi: 10.1111/j.1365-3024.2006.00772. x. PMID: 16438671.

6. Clark IA, Budd AC, Alleva LM, Cowden WB. Human malarial disease: a consequence of inflammatory cytokine release. Malar J. 2006 Oct 10; 5:85. doi: 10.1186/1475-2875-5-85. PMID: 17029647; PMCID: PMC1629020.

7. Day NP, Hien TT, Schollaardt T, Loc PP, Chuong LV, Chau TT, Mai NT, Phu NH, Sinh DX, White NJ, Ho M. The prognostic and pathophysiologic role of pro- and anti-inflammatory cytokines in severe malaria. J Infect Dis. 1999 Oct;180(4):1288–97. doi: 10.1086/315016. PMID: 10479160.

8. van der Heyde HC, Nolan J, Combes V, Gramaglia I, Grau GE. A unified hypothesis for the genesis of cerebral malaria: sequestration, inflammation and hemostasis leading to microcirculatory dysfunction. Trends Parasitol. 2006 Nov;22(11):503–8. doi: 10.1016/j.pt.2006.09.002. Epub 2006 Sep 18. PMID: 16979941.

9. Obeagu EI. Role of cytokines in immunomodulation during malaria clearance. Ann Med Surg (Lond). 2024 Apr 3;86(5):2873–2882. doi: 10.1097/MS9.0000000000002019. PMID: 38694310; PMCID: PMC11060309.

10. Gazzinelli RT, Kalantari P, Fitzgerald KA, Golenbock DT. Innate sensing of malaria parasites. Nat Rev Immunol. 2014 Nov;14(11):744–57. doi: 10.1038/nri3742. Epub 2014 Oct 17. PMID: 25324127.

11. Krishnegowda G, Hajjar AM, Zhu J, Douglass EJ, Uematsu S, Akira S, Woods AS, Gowda DC. Induction of proinflammatory responses in macrophages by the glycosylphosphatidylinositols of Plasmodium falciparum: cell signaling receptors, glycosylphosphatidylinositol (GPI) structural requirement, and regulation of GPI activity. J Biol Chem. 2005 Mar 4;280(9):860616. doi: 10.1074/jbc.M413541200. Epub 2004 Dec 28. PMID: 15623512; PMCID: PMC4984258.

12. Olivier M, Van Den Ham K, Shio MT, Kassa FA, Fougeray S. Malarial pigment hemozoin and the innate inflammatory response. Front Immunol. 2014 Feb 5; 5:25. doi: 10.3389/fimmu.2014.00025. PMID: 24550911; PMCID: PMC3913902.

13. Coban C, Ishii KJ, Kawai T, Hemmi H, Sato S, Uematsu S, Yamamoto M, Takeuchi O, Itagaki S, Kumar N, Horii T, Akira S. Toll-like receptor 9 mediates innate immune activation by the malaria pigment hemozoin. J Exp Med. 2005 Jan 3;201(1):19–25. doi: 10.1084/jem.20041836. PMID: 15630134; PMCID: PMC2212757.

14. Parroche P, Lauw FN, Goutagny N, Latz E, Monks BG, Visintin A, Halmen KA, Lamphier M, Olivier M, Bartholomeu DC, Gazzinelli RT, Golenbock DT. Malaria hemozoin is immunologically inert but radically enhances innate responses by presenting malaria DNA to Toll-like receptor 9. Proc Natl Acad Sci U S A. 2007 Feb 6;104(6):1919–24. doi: 10.1073/pnas.0608745104. Epub 2007 Jan 29. PMID: 17261807; PMCID: PMC1794278.

15. Sharma S, DeOliveira RB, Kalantari P, Parroche P, Goutagny N, Jiang Z, Chan J, Bartholomeu DC, Lauw F, Hall JP, Barber GN, Gazzinelli RT, Fitzgerald KA, Golenbock DT. Innate immune recognition of an AT-rich stem-loop DNA motif in the Plasmodium falciparum genome. Immunity. 2011 Aug 26;35(2):194–207. doi: 10.1016/j.immuni.2011.05.016. Epub 2011 Aug 4. PMID: 21820332; PMCID: PMC3162998.

16. Lamb TJ, Schenk MP, Todryk SM. How do malaria parasites activate dendritic cells? Future Microbiol. 2010 Aug;5(8):1167–71. doi: 10.2217/fmb.10.85. PMID: 20722596.

17. Palanisamy V, Jakymiw A, Van Tubergen EA, D’Silva NJ, Kirkwood KL. Control of cytokine mRNA expression by RNA-binding proteins and microRNAs. J Dent Res. 2012 Jul;91(7):651–8. doi: 10.1177/0022034512437372. Epub 2012 Feb 1. PMID: 22302144; PMCID: PMC3383846.

18. Carpenter S, Ricci EP, Mercier BC, Moore MJ, Fitzgerald KA. Post-transcriptional regulation of gene expression in innate immunity. Nat Rev Immunol. 2014 Jun;14(6):361–76. doi: 10.1038/nri3682. PMID: 24854588.

19. Turner M, Díaz-Muñoz MD. RNA-binding proteins control gene expression and cell fate in the immune system. Nat Immunol. 2018 Feb;19(2):120–129. doi: 10.1038/s41590-017-0028-4. Epub 2018 Jan 18. PMID: 29348497.

20. O’Connell RM, Rao DS, Chaudhuri AA, Baltimore D. Physiological and pathological roles for microRNAs in the immune system. Nat Rev Immunol. 2010 Feb;10(2):111–22. doi: 10.1038/nri2708. PMID: 20098459.

21. Armas P, Coux G, Weiner AMJ, Calcaterra NB. What’s new about CNBP? Divergent functions and activities for a conserved nucleic acid binding protein. Biochim Biophys Acta Gen Subj. 2021 Nov;1865(11):129996. doi: 10.1016/j.bbagen.2021.129996. Epub 2021 Aug 30. PMID: 34474118.

22. Chen Y, Sharma S, Assis PA, Jiang Z, Elling R, Olive AJ, Hang S, Bernier J, Huh JR, Sassetti CM, Knipe DM, Gazzinelli RT, Fitzgerald KA. CNBP controls IL-12 gene transcription and Th1 immunity. J Exp Med. 2018 Dec 3;215(12):3136–3150. doi: 10.1084/jem.20181031. Epub 2018 Nov 15. PMID: 30442645; PMCID: PMC6279399.

23. Gazzinelli RT, Wysocka M, Hayashi S, Denkers EY, Hieny S, Caspar P, Trinchieri G, Sher A. Parasite-induced IL-12 stimulates early IFN-gamma synthesis and resistance during acute infection with Toxoplasma gondii. J Immunol. 1994 Sep 15;153(6):2533–43. PMID: 7915739.

24. Lee E, Lee TA, Kim JH, Park A, Ra EA, Kang S, Choi HJ, Choi JL, Huh HD, Lee JE, Lee S, Park B. CNBP acts as a key transcriptional regulator of sustained expression of interleukin-6. Nucleic Acids Res. 2017 Apr 7;45(6):3280–3296. doi: 10.1093/nar/gkx071. PMID: 28168305; PMCID: PMC5389554.

25. Chen Y, Lei X, Jiang Z, Fitzgerald KA. Cellular nucleic acid-binding protein is essential for type I interferon-mediated immunity to RNA virus infection. Proc Natl Acad Sci U S A. 2021 Jun 29;118(26):e2100383118. doi: 10.1073/pnas.2100383118. PMID: 34168080; PMCID: PMC8255963.

26. Stevenson MM, Tam MF, Wolf SF, Sher A. IL-12-induced protection against blood-stage Plasmodium chabaudi AS requires IFN-gamma and TNF-alpha and occurs via a nitric oxide-dependent mechanism. J Immunol. 1995 Sep 1;155(5):2545–56. PMID: 7650384.

27. Yoshimoto T, Yoneto T, Waki S, Nariuchi H. Interleukin-12-dependent mechanisms in the clearance of blood-stage murine malaria parasite Plasmodium berghei XAT, an attenuated variant of P. berghei NK65. J Infect Dis. 1998 Jun;177(6):1674–81. doi: 10.1086/515301. PMID: 9607848.

28. Su Z, Stevenson MM. Central role of endogenous gamma interferon in protective immunity against blood-stage Plasmodium chabaudi AS infection. Infect Immun. 2000 Aug;68(8):4399–406. doi: 10.1128/IAI.68.8.4399-4406.2000. PMID: 10899836; PMCID: PMC98333.

29. Chen Y, Lei X, Jiang Z, Humphries F, Parsi KM, Mustone NJ, Ramos I, Mutetwa T, Fernandez-Sesma A, Maehr R, Caffrey DR, Fitzgerald KA. Cellular nucleic acid-binding protein restricts SARS-CoV-2 by regulating interferon and disrupting RNA-protein condensates. Proc Natl Acad Sci U S A. 2023 Nov 21;120(47):e2308355120. doi: 10.1073/pnas.2308355120. Epub 2023 Nov 14. PMID: 37963251; PMCID: PMC10666094.

30. Del Portillo HA, Ferrer M, Brugat T, Martin-Jaular L, Langhorne J, Lacerda MV. The role of the spleen in malaria. Cell Microbiol. 2012 Mar;14(3):343–55. doi: 10.1111/j.1462-5822.2011.01741.x. Epub 2012 Feb 2. PMID: 22188297.

31. Stevenson MM, Riley EM. Innate immunity to malaria. Nat Rev Immunol. 2004 Mar;4(3):169–80. doi: 10.1038/nri1311. PMID: 15039754.

32. Popa GL, Popa MI. Recent Advances in Understanding the Inflammatory Response in Malaria: A Review of the Dual Role of Cytokines. J Immunol Res. 2021 Nov 8;2021:7785180. doi: 10.1155/2021/7785180. PMID: 34790829; PMCID: PMC8592744.

33. Cai C, Hu Z, Yu X. Accelerator or Brake: Immune Regulators in Malaria. Front Cell Infect Microbiol. 2020 Dec 10;10:610121. doi: 10.3389/fcimb.2020.610121. PMID: 33363057; PMCID: PMC7758250.

34. Orengo JM, Leliwa-Sytek A, Evans JE, Evans B, van de Hoef D, Nyako M, Day K, Rodriguez A. Uric acid is a mediator of the Plasmodium falciparum-induced inflammatory response. PLoS One. 2009;4(4):e5194. doi: 10.1371/journal.pone.0005194. Epub 2009 Apr 17. PMID: 19381275; PMCID: PMC2667251.

35. de Souza JB, Hafalla JC, Riley EM, Couper KN. Cerebral malaria: why experimental murine models are required to understand the pathogenesis of disease. Parasitology. 2010 Apr;137(5):755–72. doi: 10.1017/S0031182009991715. Epub 2009 Dec 23. PMID: 20028608.

36. Grau GE, Kindler V, Piguet PF, Lambert PH, Vassalli P. Prevention of experimental cerebral malaria by anticytokine antibodies. Interleukin 3 and granulocyte macrophage colony-stimulating factor are intermediates in increased tumor necrosis factor production and macrophage accumulation. J Exp Med. 1988 Oct 1;168(4):1499–504. doi: 10.1084/jem.168.4.1499. PMID: 3049913; PMCID: PMC2189068.

37. Buffet PA, Safeukui I, Milon G, Mercereau-Puijalon O, David PH. Retention of erythrocytes in the spleen: a double-edged process in human malaria. Curr Opin Hematol. 2009 May;16(3):157–64. doi: 10.1097/MOH.0b013e32832a1d4b. PMID: 19384231.

38. Schroder K, Hertzog PJ, Ravasi T, Hume DA. Interferon-gamma: an overview of signals, mechanisms and functions. J Leukoc Biol. 2004 Feb;75(2):163–89. doi: 10.1189/jlb.0603252. Epub 2003 Oct 2. PMID: 14525967.

39. Su Z, Stevenson MM. IL-12 is required for antibody-mediated protective immunity against blood-stage Plasmodium chabaudi AS malaria infection in mice. J Immunol. 2002 Feb 1;168(3):1348–55. doi: 10.4049/jimmunol.168.3.1348. PMID: 11801675.

40. Curion F, Handel AE, Attar M, Gallone G, Bowden R, Cader MZ, Clark MB. Targeted RNA sequencing enhances gene expression profiling of ultra-low input samples. RNA Biol. 2020 Dec;17(12):1741–1753. doi: 10.1080/15476286.2020.1777768. Epub 2020 Jun 28. PMID: 32597303; PMCID: PMC7746246.

41. Riley EM, Stewart VA. Immune mechanisms in malaria: new insights in vaccine development. Nat Med. 2013 Feb;19(2):168–78. doi: 10.1038/nm.3083. PMID: 23389617.

42. de Souza Silva L, Monks BG, Forconi CS, Crabtree JN, De Paula Tamburro N, Kurt-Jones EA, Gazzinelli RT, Fitzgerald KA, Golenbock DT. Interleukin-10 limits immune-mediated pathology in chronic subclinical plasmodial infection. PLoS Negl Trop Dis. 2025 Sep 19;19(9):e0013554. doi: 10.1371/journal.pntd.0013554. PMID: 40972121; PMCID: PMC12463326.

43. https://support-docs.illumina.com/LP/IlluminaStrandedmRNA/Content/LP/FrontPages/IlluminaStrandedmRNA.htm

44. Langmead B, Salzberg SL. Fast gapped-read alignment with Bowtie 2. Nat Methods. 2012 Mar 4;9(4):357–9. doi: 10.1038/nmeth.1923. PMID: 22388286; PMCID: PMC3322381.

45. Li B, Dewey CN. RSEM: accurate transcript quantification from RNA-Seq data with or without a reference genome. BMC Bioinformatics. 2011 Aug 4;12:323. doi: 10.1186/1471-2105-12-323. PMID: 21816040; PMCID: PMC3163565.

46. Leng N, Li Y, McIntosh BE, Nguyen BK, Duffin B, Tian S, Thomson JA, Dewey CN, Stewart R, Kendziorski C. EBSeq-HMM: a Bayesian approach for identifying gene-expression changes in ordered RNA-seq experiments. Bioinformatics. 2015 Aug 15;31(16):2614–22. doi: 10.1093/bioinformatics/btv193. Epub 2015 Apr 5. PMID: 25847007; PMCID: PMC4528625.

47. Falcon S, Gentleman R. Using GOstats to test gene lists for GO term association. Bioinformatics. 2007 Jan 15;23(2):257–8. doi: 10.1093/bioinformatics/btl567. Epub 2006 Nov 10. PMID: 17098774.

48. R Core Team (2024). _R: A Language and Environment for Statistical Computing_. R Foundation for Statistical Computing, Vienna, Austria. https://www.R-project.org/

49. Wickham H (2016). ggplot2: Elegant Graphics for Data Analysis. Springer-Verlag New York. ISBN 978-3-319-24277-4, https://ggplot2.tidyverse.org

